# Hairpin-Functionalized Gold Nanoparticles as an Adaptable Platform for Detecting MicroRNA Signatures

**DOI:** 10.64898/2025.12.04.692125

**Authors:** Catarina Coutinho, Nuria Lafuente-Gómez, Irene de la Iglesia, Luis A. Campos, Demián Pardo, Irene Pardo, Jorge Royes, Rebeca Bocanegra, Milagros Castellanos, Álvaro Somoza

**Affiliations:** Instituto Madrileño de Estudios Avanzados en Nanociencia (IMDEA Nanociencia) Faraday 9, Ciudad Universitaria de Cantoblanco, 28049 Madrid, Spain

**Keywords:** gold nanoparticle-based sensors, microRNAs, nucleic acid signatures, molecular profiling, liquid biopsy

## Abstract

Early, accurate, and fast diagnosis is essential to ensure positive health outcomes, through effective treatment interventions and disease control, as well as the study of physiological changes related to gene expression. Liquid biopsies and point-of-care (PoC) detection are valuable tools to achieve this goal, allowing the study and monitoring of a patients’ molecular profile in a timely and simple manner. MicroRNAs (or miRNAs) have been proposed as biomarkers for detection in liquid biopsies, as they are stable in body fluids and their dysregulation is associated with many diseases. In this work, a sensor based on gold nanoparticles (AuNPs) functionalized with oligonucleotides is reported, aiming for the detection of nucleic acids, in particular miRNAs. This sensor is based on the recognition of target sequences by hairpin-shaped oligonucleotide probes, which allows the modulation of the colloidal stability of the AuNPs, producing color changes detectable with the naked eye. The system is used to detect a panel of miRNAs, demonstrating its versatility for the detection of relevant nucleic acid signatures. The sensor detects single miRNAs with good sensitivity and selectivity and, what is more, it can be used to recognize several miRNAs simultaneously at picomolar concentrations. The system was further adapted to a lateral flow assay (LFA) format, producing a visible colored line on lateral flow test strips, and coupled with isothermal amplification to reach femtomolar detection levels. Its properties make it suitable for use in the point-of-care (PoC), contributing to fast and early detection of pathological and physiological molecular profiles.

## 1. Introduction

Accurate, rapid, and early disease diagnosis is fundamental for enabling timely therapeutic interventions and improving clinical outcomes.^1,2^ In this context, the detection of overexpressed molecular markers linked to gene expression changes, whether resulting from pathological dysregulation or from physiological activation processes, has become increasingly important.^3^ Such molecular insights not only enhance diagnostic precision, but also support the prevention and control of infectious diseases, and enable more effective monitoring of treatment responses.^1,3^ Over the past years, non-invasive methods like liquid biopsy have gradually replaced invasive techniques in disease diagnosis and monitoring.^4^ It consists of the detection and monitoring of overexpressed biomarkers in different body fluids. These biomarkers include nucleic acids (e.g., ctDNAs, miRNAs), proteins, exosomes, and cells.^5^ Apart from being non-invasive, liquid biopsies allow fast and accurate analyses, early detection, and repeated sample collection with simple procedures, in some cases, even by the patients themselves. This enables both sample collection and detection in the point-of-care (PoC), as well as the study and monitoring of a patients’ molecular profile, allowing for early, accurate, and fast diagnosis, and to study physiological processes.^4–6^

MicroRNAs (or miRNAs), in particular, have been widely studied as candidate biomarkers for detection in liquid biopsies.^7^ MiRNAs are a family of non-coding RNAs that regulate gene expression by binding to messenger RNAs (mRNAs), promoting their cleavage or translational repression.^8,9^ MiRNAs regulate essential processes like development, differentiation, proliferation, homeostasis, metabolism, and response to infection. Conversely, miRNAs are dysregulated in various diseases, including cancer, cardiovascular and metabolic diseases, viral and bacterial infections, and nervous system disorders.^10–12^ MicroRNAs have been detected in fluids such as plasma, serum, urine, saliva, seminal and ascites fluids, amniotic pleural effusions, or cerebrospinal fluid.^13–18^

Despite their convenient use as non-invasive biomarkers, the detection of these nucleic acids presents some difficulties, and their general use in clinics is still limited.^19^ It is also important to note that, in most cases, changes in the expression of a single miRNA are not enough for the clear identification of diseases or physiological changes. For this reason, profiling of the expression of several miRNAs is generally used to characterize disease or physiological states.^20^ Regarding miRNA detection, conventional methods like RT-qPCR, microarray analysis, Northern blotting, polyacrylamide gel electrophoresis, or the more recent microarray and RNA-Seq platforms offer high accuracy and sensitivity.^19,21^ However, these methods are not suitable for routine diagnosis because, first, highly trained personnel and expensive equipment are required, and, second, they are time-consuming.^22^ For these reasons, the development of new detection systems, particularly those aiming for liquid biopsy detection in the point-of-care (PoC), is very welcome. In this regard, the use of nanomaterials, in particular, gold nanoparticles (AuNPs), has eased the development of this kind of sensing devices. AuNPs have unique and valuable properties for biosensor design, including high surface-to-volume ratio, high surface energy and reactivity, tunable optical properties, high stability, and biocompatibility.^23,24^ These features have improved the applicability of sensor systems in PoC, in terms of simplicity, sensitivity, selectivity, cost-effectiveness, and detection speed.^25^ Colorimetric sensors in solution and lateral flow assays (LFAs) are explored in this work, two systems that meet these PoC characteristics due to their simple readouts, based on color changes.

For the detection of nucleic acids, in particular, AuNPs have been employed extensively due to their excellent optical properties and ease of preparation and modification.^26^ The high extinction coefficient of AuNPs, and the ability to modulate their color depending on shape, size, and aggregation state, have motivated their extensive use in colorimetric sensors, both in solution and LFAs.^22,26^ In most cases, nucleic acid detection involves the functionalization of AuNPs with complementary nucleic acid probes. The first AuNP-based nucleic acid sensors were proposed by Prof. C. Mirkin *et al.*, and consisted of an AuNP core densely functionalized with linear DNA oligonucleotides, complementary to a target of interest.^27–30^

In this regard, we have prepared AuNPs modified with hairpin-shaped oligonucleotides, which present a stable folded structure that unfolds only in the presence of a complementary target sequence. The structure’s opening exposes a molecule that can produce a signal. For instance, a fluorescent dye at the end of the oligonucleotide can be used as a reporter molecule, constituting the commonly named molecular beacons.^31^ In this case, the oligonucleotides, once conjugated to AuNPs, can be used for intracellular detection of RNA. In this system, the unfolding of the structure, upon nucleic acid hybridization, leads to an increase in fluorescence.^32^

In our case, we use oligonucleotides that fold into a hairpin structure (molecular beacon-like oligonucleotides, MBs), and contain a hydrophobic molecule (a cholesterol derivative) at one end, and a thiol group at the other, used to functionalize AuNPs (**Figure 1A**). The obtained nanostructure exhibits an initial arrangement in which the hairpin retains the cholesterol derivative buried inside it.^33,34^ In this configuration, the nanoparticle is well dispersed in aqueous media, and presents the characteristic reddish color due to the plasmon band of AuNPs.^35^ However, in the presence of the specific target sequence, the hairpin unfolds, exposing the cholesterol to the water molecules. This rearrangement induces the aggregation of the structure, which can be detected by the naked eye – colorimetric sensor in solution (**Figure 1B**). We further adapted this system to a LFA format, replacing the terminal cholesterol moieties with biotin moieties in the hairpin oligonucleotides. These molecules remain inside the nanostructure, but are exposed in the presence of the target sequence.^36,37^ The mixture is then loaded onto LFA strips containing a streptavidin test line, leading to the retention of the AuNPs exposing biotin derivatives, due to the high affinity between the two molecules, and producing a visible colored line (**Figure 1C**).

**Figure 1.**
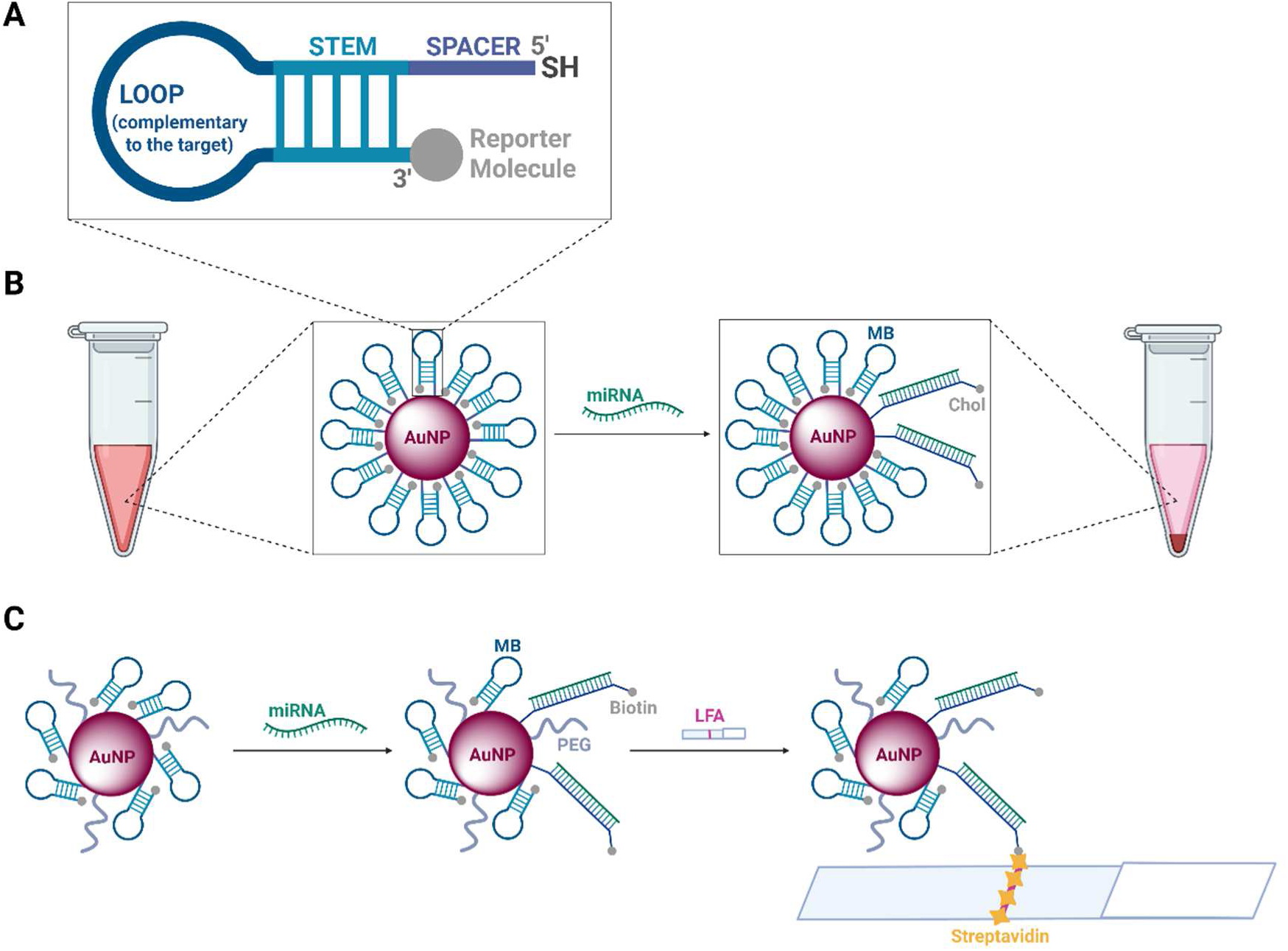
Schematic representation of the proposed miRNA sensors, and the molecular mechanisms involved in the detection process. (**A**) Structure of the molecular beacon-like probes (MBs) used in this work. The oligonucleotides contain a thiol moiety (SH) at the 5’-end, and a reporter molecule (cholesterol or biotin) at the 3’-end. The central sequence of the oligonucleotide (loop) is complementary to the target miRNA, and is flanked by additional sequences that recognize each other (stem). Therefore, the oligonucleotide can fold, yielding a stable hairpin structure. A spacer of 5 thymines (T) is also included between the stem sequence and the thiol moiety, used to functionalize the AuNPs. (**B**) Colorimetric sensors (in solution) – In the presence of the target miRNA, the hairpin structure unfolds, exposing the cholesterol moiety, and leading to the aggregation and precipitation of the AuNPs. (**C**) LFA sensors - Binding of the target miRNA leads to opening of the hairpin structure and exposure of the biotin molecule. Loading onto streptavidin lateral flow strips leads to retention of the AuNPs on the test line, producing a colored signal. Image created with BioRender.com

Here, we report the use of these approaches for the detection of a combination of miRNAs, developing multiplexed systems that allow visual detection of miRNAs in the nanomolar to picomolar range. These systems were further coupled with isothermal amplification, in particular, the Exponential Amplification Reaction (EXPAR), to reach femtomolar detection levels. The EXPAR technique allows amplification of short nucleic acids with high efficiency, and has been extensively used in miRNA detection.^38–42^ We believe that the development of user friendly and accessible detection devices will help clinicians establish a quick and precise diagnosis, and prognosis, of different diseases, with measurable nucleic acid biomarkers in body fluids. These biomarkers can be further used to study physiological changes related to gene expression and activation pathways. As a model system to simulate a panel of miRNAs, we study the detection of the miRNAs 146a, 155, and 191, which are related with inflammation, immune response, cancer, and other diseases.^43–47^

## 2. Material and Methods

### 2.1. Oligonucleotides

The oligonucleotides used in this work were either purchased or synthesized in the laboratory. The oligonucleotide providers were Integrated DNA Technologies (IDT, Coralville, IA, USA), Sigma Aldrich (San Luis, MO, USA), or Biomers GmbH (Ulm, Germany). Oligonucleotides (DNA-based) were also synthesized by solid-phase synthesis using phosphoramidites (LGC Biosearch Technologies (Steinach, Germany) and Wuhu Huaren (Wuhu, Anhui, China)), in a H8/H6 DNA/RNA synthesizer (K&A Labs GmbH, Schaafheim, Germany), as described previously with some modifications.^32^

After the synthesis, the solid supports were transferred to screw-cap glass vials and incubated overnight with 2 mL of a 32% ammonia solution (Sigma Aldrich). Then, the supernatants (containing the oligonucleotide separated from the solid support) were transferred to 2 mL microcentrifuge tubes and dried in a centrifugal evaporator (Eppendorf, Hamburg, Germany). The oligonucleotides were then resuspended and purified by 20% polyacrylamide gel electrophoresis, followed by electroelution from gel fractions into dialysis tubes (3.5 KDa cut-off, Spectrum, ThermoFisher Scientific, Waltham, MA, USA). Finally, the samples were desalted using Amersham NAP-10 Columns (Cytiva, Marlborough, MA, USA) and dried in the centrifugal evaporator.

Both commercial and synthesized oligonucleotides were resuspended in water and quantified by UV-Vis spectra (λ = 260 nm) in a Cary 5000 UV-Vis-NIR spectrophotometer (Agilent, Santa Clara, CA, USA). The OligoAnalyzer tool from IDT was used to determine the molar extinction coefficient.^48^ After quantification, the oligonucleotide solutions were aliquoted and stored at -20 °C until further use.

In this work, hairpin-shaped DNA molecules that unfold in the presence of a complementary sequence, similar to molecular beacons^31^ (molecular beacon-like oligonucleotides, MBs), were used as probes for nucleic acid detection. These oligonucleotides were modified with a thiol at one end, and either a cholesterol or a biotin derivative at the other. Their sequences are listed in **Table 1**, along with their targets and other non-target sequences (scrambles).

**Table 1.**
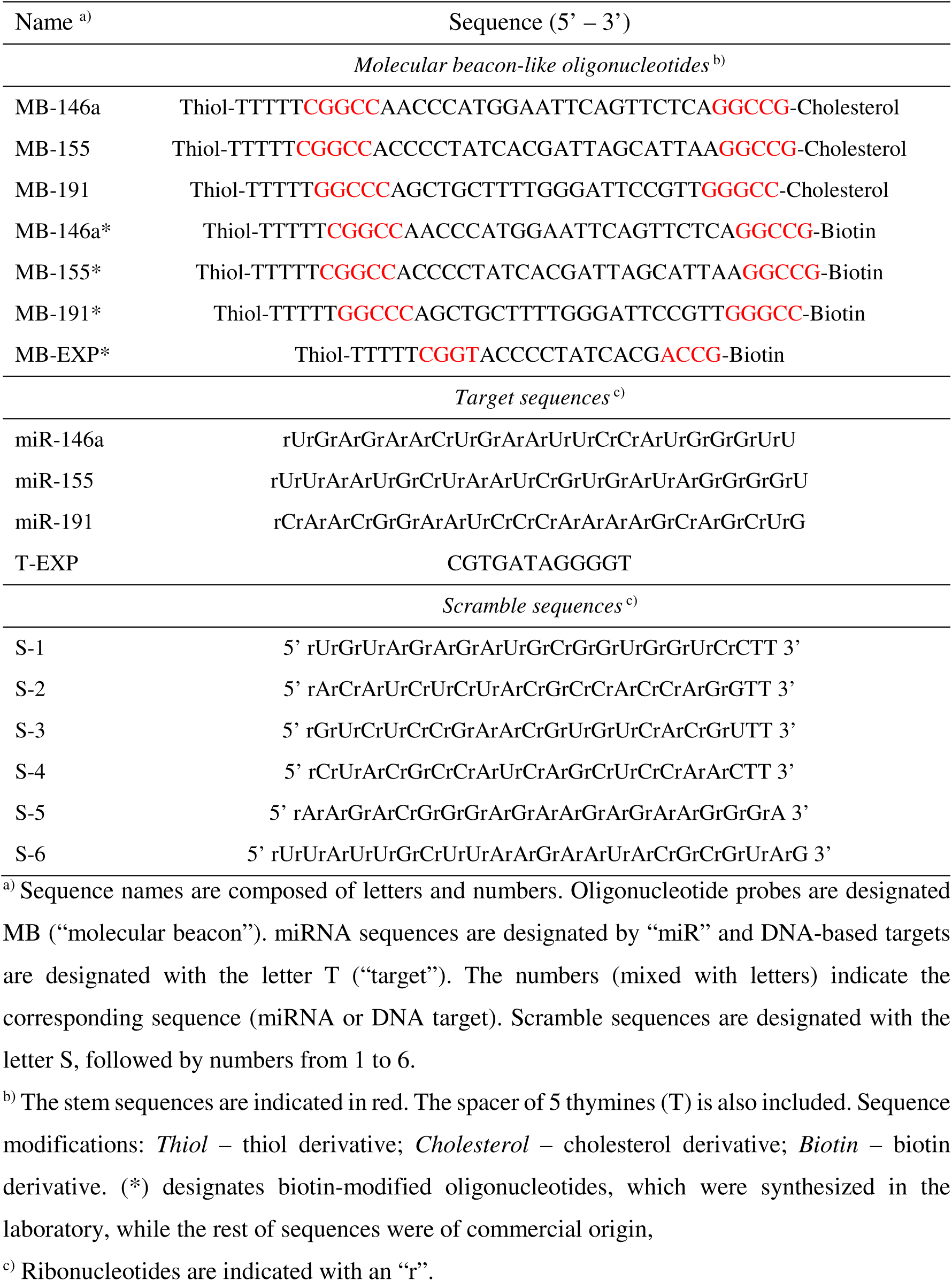
Oligonucleotide sequences employed in this work.

### 2.2. Synthesis of AuNPs

AuNPs of 40 nm were purchased from NanoImmunoTech (Vigo, Spain). 12 nm AuNPs were synthesized in the laboratory by the Turkevich-Frens method,^49,50^ by diluting 77 µL of a 1.23 M solution of chloroauric acid (HAuCl_4_) (ThermoFisher Scientific) in 100 mL of boiling water, under reflux and stirring. Then, 10 mL of a 0.04 M solution of trisodium citrate was added to the solution and incubated for 15 minutes. After overnight stirring at room temperature (RT), the AuNP solution was filtered using a 20-40 µm glass filter plate under vacuum, followed by a 0.22 µm polyethersulfone (PES) syringe filter. The concentration of both synthesized and commercial AuNPs was determined from their absorbance spectra, according to Haiss *et al.*,^51^ using the values at their absorbance plasmon peak and those at λ = 450 nm, measured in the Cary 5000 spectrophotometer.

### 2.3. Preparation of AuNP-based sensors

For colorimetric sensors (in solution), 12 nm AuNPs were used at 7 nM concentration and combined with 1.33 pmol of cholesterol-modified MBs per µ L of AuNP solution, in order to achieve maximum surface coverage, with a two-fold excess.^34^ Before functionalization, the MBs were treated with tris(2-carboxyethyl) phosphine hydrochloride (TCEP) (Sigma Aldrich) to remove the protecting groups from the thiol modifications.^34,52^ Then, the deprotected MBs were added to the AuNP solution, followed by NaCl addition, gradually to 0.3 M, to increase MB binding.^29,53,54^ The obtained AuNPs were incubated at RT overnight and then washed by iterative centrifugation (Eppendorf 5427R centrifuge), using Milli-Q^®^ water (Merck Millipore, Burlington, MA, USA). The supernatants resulting from the first washing step were collected and used to measure the loading efficiency of each AuNP formulation, by quantification of the unbound MB, using UV-Vis absorbance (λ = 260 nm) in a Synergy H4 microplate reader (BioTek, Winooski, VT, USA).^34^

For lateral flow-based sensors, 40 nm AuNPs were used at 2 nM concentration and combined with 2.23 pmol of biotin-modified MBs per µ L of AuNP solution.^34^ The MBs were also treated with TCEP prior to functionalization as described above. After that, the deprotected MBs were added to the AuNP solution, followed by overnight incubation at RT. The next day, 1.3 µ L of lipoic acid-modified polyethylene glycol (LP-PEG) of 3 KDa was added and incubated for 1 hour. This molecule was synthesized as described in previous work,^52,55^ (see Supporting Information, *Section 1*), and used to stabilize the nanostructure, since in this case NaCl additions were not performed, in order to avoid the non-specific unfolding of the biotin-containing MBs due to a high loading. After incubation, the AuNPs were washed and their loading efficiency was determined as described above.

### 2.3. Characterization of AuNPs

AuNPs were visualized by transmission electron microscopy (TEM) using a 120 KeV JEM1400 Flash (Jeol, Akishima, Tokyo, Japan) at the CBMSO-CSIC/UAM institute (Madrid, Spain). Samples were prepared by placing a glow-discharged carbon-shadowed formvar-coated HEX 400-mesh copper grid (Agar Scientific, Stansted, Essex, UK) over one drop of the AuNP solution for 2 minutes and drying the excess of sample. The shape and size distributions were determined through an automated analysis of randomly selected areas in the TEM images, using the Analyze Particles function of ImageJ software. The resulting size histograms were fitted to Gaussian distributions using the Excel software. The mean hydrodynamic diameter and Zeta potential of the AuNPs were determined by dynamic light scattering (DLS) and electrophoretic light scattering (ELS), respectively, using a Zetasizer Nano ZS instrument (Malvern Panalytical, Malvern, UK). For simple optical characterization purposes (not quantification), the UV-Vis absorbance spectra were collected in the Synergy H4 microplate reader.

### 2.4. Preparation of streptavidin lateral flow strips

Nitrocellulose (NC) membranes (Whatman FF80HP, Cytiva) of 4 cm width were used for the preparation of lateral flow strips. An Automated Lateral Flow Reagent Dispenser (ALFRD, Claremont BioSolutions, Upland, CA, USA), coupled to a Fusion 200 Syringe Pump (Chemyx, Stafford, TX, USA), was used to deposit all the reagents onto the NC membranes. The immobilization of streptavidin onto the NC membranes followed the method proposed by Másson *et al.*, using diaminoalkanes and glutaraldehyde (GA).^56^ First, a 5% solution of 1,4-diaminobutane (Sigma Aldrich) in water was deposited on the NC membranes and incubated at RT for 1 hour. Then, the NC membranes were washed twice with water and dried at RT for 16 hours. The next day, a 0.1% solution of GA (Sigma Aldrich) in 0.5 M carbonate-bicarbonate buffer (Na_2_CO_3_/NaHCO_3_, pH=10) was prepared and filtered with a 0.2 µm polyvinylidene difluoride (PVDF) syringe filter. The filtered solution was deposited over the 1,4-diaminobutane line on the NC membranes and incubated at RT for 15 minutes, protected from light. After that, the NC membranes were washed twice and dried as in the previous step. Finally, a 1 mg/mL solution of streptavidin from *Streptomyces avidinii* (Sigma Aldrich) in water was deposited over the previous lines and incubated at RT for 6 hours. Then, the NC membranes were washed twice in 10 mM Tris-HCl buffer (pH=8) with 0.1% Tween-20 and dried as previously described.

After immobilization, the NC membranes were mounted onto adhesive backing cards (DCN Dx, Carlsbad, CA, USA), and C083 cellulose fiber sample/absorbent pads (Merck Millipore) were placed adjacent to the NC membranes, with a 2 mm overlap. Finally, the mounted NC membranes were cut into 0.5 cm width NC strips, using a rotary trimmer, and stored at 4 °C for further use.

### 2.5. MiRNA detection assays

The detection of miRNAs in solution was performed as previously reported.^33,34^ Briefly, 50 µ L of 12 nm AuNPs functionalized with cholesterol-modified MBs were mixed with 5 µ L of target sequences (at different concentrations), 5 µ L of NaCl 5 M, and 100 µ L of PBS 1X (ThermoFisher Scientific), for a constant final volume of 160 µ L, in 1.5 mL SuperClear^®^ microcentrifuge tubes (Labcon, Petaluma, CA, USA). Two sedimentation procedures were studied: gravity-dependent or assisted by low-speed centrifugation at 5000 rpm for 5 minutes. For absorbance measurements, 70 µL from the upper part of each sample were removed slowly, and transferred to a UV-Vis absorbance 96-well microplate (Corning, NY, USA). The spectra from 400 to 650 nm were collected using the Synergy H4 microplate reader, and the absorbance maxima were determined. Experiments were done in triplicates, and data were collected at different time points, as well as photographs of the samples.

For lateral flow-based detection, 10 µ L of 40 nm AuNPs functionalized with biotin-modified MBs were incubated (RT, 20 minutes) with 2.5 µ L of target sequences (at different concentrations), 4 µL of running buffer concentrated 5X, and 3.5 µL of nuclease-free water, for a constant final volume of 20 µ L, in 1.5 mL SuperClear^®^ microcentrifuge tubes (Labcon). The composition of the running buffer was 100 mM Phosphate Buffer, 600 mM NaCl, 4% BSA, 0.1% Tween-20. A 5X concentrated version was prepared for use in lateral flow assays. The NC strips were rehydrated with 20 µ L of 100 mM Phosphate Buffer and, after the 20-minute incubation, the AuNP mixtures were loaded onto the strips. After the samples flowed till the end of the strips, two washing steps with 20 µ L of water were performed. Then, the strips were left to dry at RT for 15-30 minutes, and imaged in a ChemiDoc Imaging System (BioRad Laboratories, Hercules, CA, USA). The signal intensities were quantified from the obtained images using the ImageJ software.

### 2.6. EXPAR amplification and detection of miRNAs

The amplification of the miR-155 sequence was performed by the Exponential Amplification Reaction (EXPAR).^38–42^ The oligonucleotides used in this reaction included a specifically designed DNA template (Temp), and a DNA-based analog of miR-155 used as target (T-155). Oligonucleotides were purchased from Sigma Aldrich and IDT, while all the remaining reagents were purchased from New England Biolabs (Ipswich, MA, USA), except SYBR Green I, which was obtained from ITW Reagents (Monza, Italy). Oligonucleotide sequences are listed in **Table 2**.

**Table 2.**
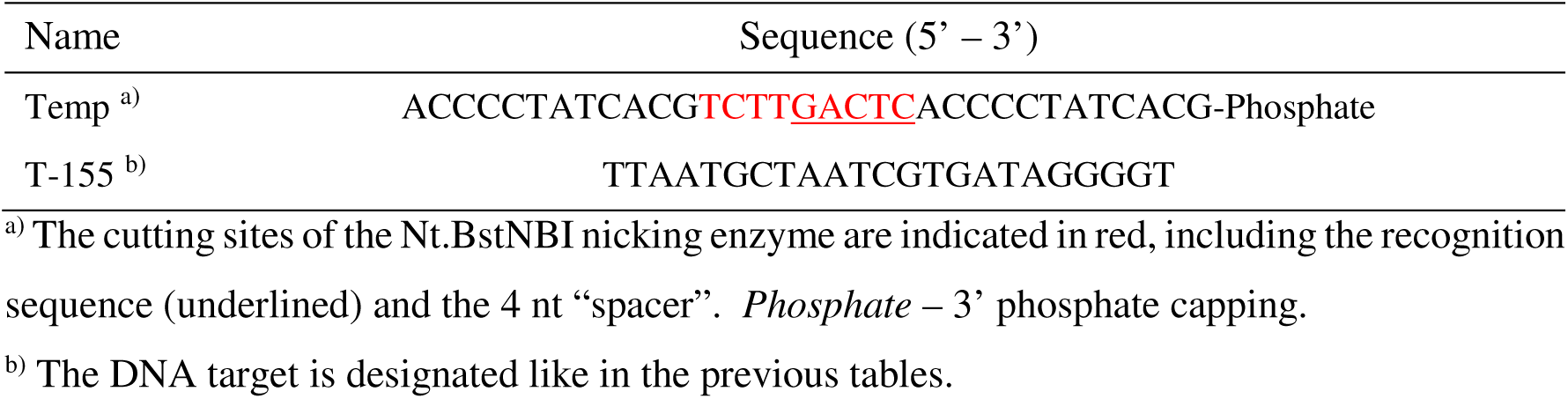
Oligonucleotides used for the EXPAR-based amplification of miR-155.

The EXPAR reactions were performed according to reported methods, with some modifications.^38–42^ The reaction mixtures were prepared in two parts. Part A contained 2X SYBR Green I, 200 nM DNA template (Temp), 500 µM dNTPs, and T-155 at the desired concentration. Part B contained 2X ThermoPol^®^ Buffer, 1X NEBuffer 3.1, 0.16 U/µ L Vent (exo-) DNA polymerase, and 0.8 U/µ L Nt.BstNBI nicking endonuclease. Both parts were prepared on ice, in microcentrifuge tubes, and transferred separately to the wells of a white 96-well qPCR plate (half-skirted, ABI-type, notch A12) (VWR, Radnor, PA, USA). The plate was heated to 51 °C for 5 minutes in a Thermal Shaker *lite* (VWR). Then, equal volumes of the two parts were combined using a multichannel pipette, meaning that the concentration of each reagent was half the indicated, for a final volume of 20 µ L. Finally, the plate was transferred to a qTOWER^3^ G Real-Time PCR Thermal Cycler (Analytik Jena, Jena, Germany), which was also pre-heated at 51 °C, and the reaction was monitored in real-time with fluorescence measurements every 30 s.

For lateral flow-based detection of the EXPAR amplification product, parts A and B were prepared on ice in PCR tubes, and then heated to 51 °C in the Thermal Shaker. Then, the two parts were mixed for 20-µ L final reaction volumes, and incubated at 51 °C. The reactions were stopped at the desired time by rapidly placing the tubes on ice. Finally, the samples were used as target solutions in lateral flow detection assays, as described above. The detection mixture consisted in 1 µ L of EXPAR sample (without purification), 10 µ L of AuNPs functionalized to detect the EXPAR product, 4 µ L of running buffer concentrated 5X, and 5 µ L of water, for a constant final volume of 20 µ L.

### 2.7. Statistical analysis

Statistical analyses were performed using one-away analysis of variance (ANOVA) in order to compare multiple groups. Differences between groups were assessed using Tukey’s *post hoc* test. Differences were considered significant at p < 0.1. Statistical significance was denoted if the p-value was lower than 0.1 by · (p<0.1), 0.05 by * (p < 0.05), 0.01 by ** (p < 0.01), and 0.001 by *** (p < 0.001). The analyses were performed using “R” software program.^57^

## 3. Results and Discussion

### 3.1. AuNP-based colorimetric sensors in solution

#### 3.1.1. Preparation and characterization of the sensors

For the preparation of colorimetric sensors (in solution), citrate-stabilized AuNPs of 12 nm were prepared following the Turkevich-Frens approach,^49,50^ and modified with oligonucleotides that contained a thiol moiety at one end and a cholesterol derivative at the other, folding into a hairpin structure, as mentioned above (**Figure 1B**).

The obtained citrate-stabilized AuNPs were imaged by TEM, mostly showing the expected spherical morphology (**Figure 2A**). Using the Analyze Particles function of ImageJ software, histograms of the AuNP diameters were obtained and were fitted to a Gaussian curve, in Excel, to determine the mean diameter. The AuNPs presented a narrow size distribution, with a mean diameter of 12.3 ± 0.6 nm (**Figure 2B**).

**Figure 2.**
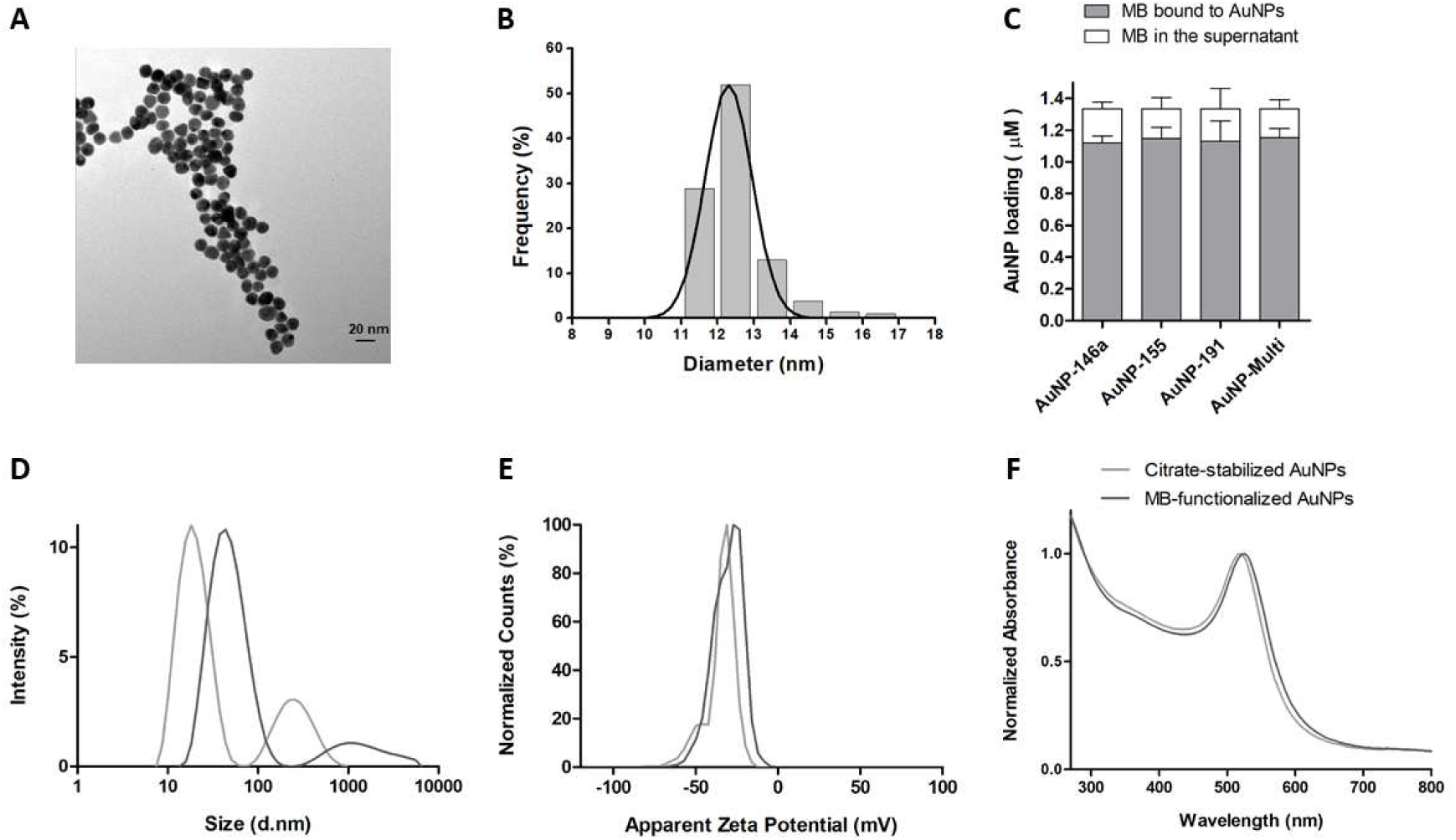
Characterization of AuNP-based sensors in solution. (**A**) TEM image of 12 nm citrate-stabilized AuNPs. (**B**) Size distribution obtained from TEM images, by measuring at least 200 AuNPs, using the Analyze Particles function of ImageJ software, and fitting to a Gaussian curve in Excel program. (**C**) Loading efficiency of 12 nm AuNPs with cholesterol-modified MBs. Results are presented as Mean ± SD: n=5 for AuNP-146a and AuNP-191; n=4 for AuNP-155; and n=6 for AuNP-Multi. (**D**) Hydrodynamic diameter, (**E**) Zeta potential distribution, and (**F**) UV-Vis absorbance spectra of both citrate-stabilized (light grey) and MB-functionalized AuNPs (dark grey). Data are shown for AuNP-191, representative of all the nanostructures, which had similar features.

The AuNPs were modified with the oligonucleotides MB-146a, MB-155, and MB-191, aimed respectively to detect the sequences miR-146a, miR-155, and miR-191 (**Table 1**). The resulting nanostructures were designated AuNP-146a, AuNP-155, and AuNP-191. Additionally, a nanostructure functionalized with an equal mixture of the three MBs was prepared for multiplexed detection and was designated AuNP-Multi. The loading efficiency of each nanostructure was determined by quantification of the MBs remaining in the supernatant from the first washing step (**Figure 2C**). All the nanostructures presented high loading efficiencies of around 85%, confirming that the oligonucleotides were added in excess, and that a significant amount of MBs were attached to the AuNPs.

The AuNPs were also characterized before and after functionalization by DLS, ELS, and UV-Vis spectroscopy. The hydrodynamic diameter (**Figure 2D**) of the nanoparticles increased from around 20 nm to 50 nm, which was reasonable to account for the influence on the hydrodynamic diameter of the presence of the MBs.^58,59^ Before and after functionalization, the AuNPs had a polydispersity index (PDI) of 0.3, influenced by presence of some nanoparticle aggregates, which were tolerated for the development of the sensors. The Zeta potential (**Figure 2E**) remained unchanged at around -35 mV, which was expected due to the exchange of the citrate molecules with the MBs, both negatively charged. Finally, the absorbance spectra (**Figure 2F**) of functionalized AuNPs were similar to the ones of bare AuNPs, highlighting the overall stability of the nanostructures. A slight red shift of the plasmon band, from 520 nm to 525 nm was observed, pointing also to the presence of the MBs on the surface of the AuNPs.^60,61^

#### 3.1.2. Sensor performance: sensitivity and selectivity

The sensors developed herein are based on color changes resulting from the aggregation and precipitation of the AuNPs out of the solution, due to the opening of the MB structure upon target binding and exposure of the cholesterol moieties.^33,34^ These properties were verified when incubating the sensors with 250 nM of their target. The absorbance of the samples (**Figure 3A**) decreased significantly after incubation with the target, to values of 50-60% of that of the blank samples, consistent with the precipitation of the AuNPs. In addition, the absorbance maxima were shifted to longer wavelengths and the bandwidth increased, which was also related to the formation of bigger particles and aggregates.^62^ No precipitation or color changes were observed when the target was not present (blank sample), while the precipitation of the particles could be detected clearly with the naked eye when incubated with the target, as a darker precipitate at the bottom of the tube, as well as the loss of the reddish color of the solution (**Figure 3B**).

**Figure 3.**
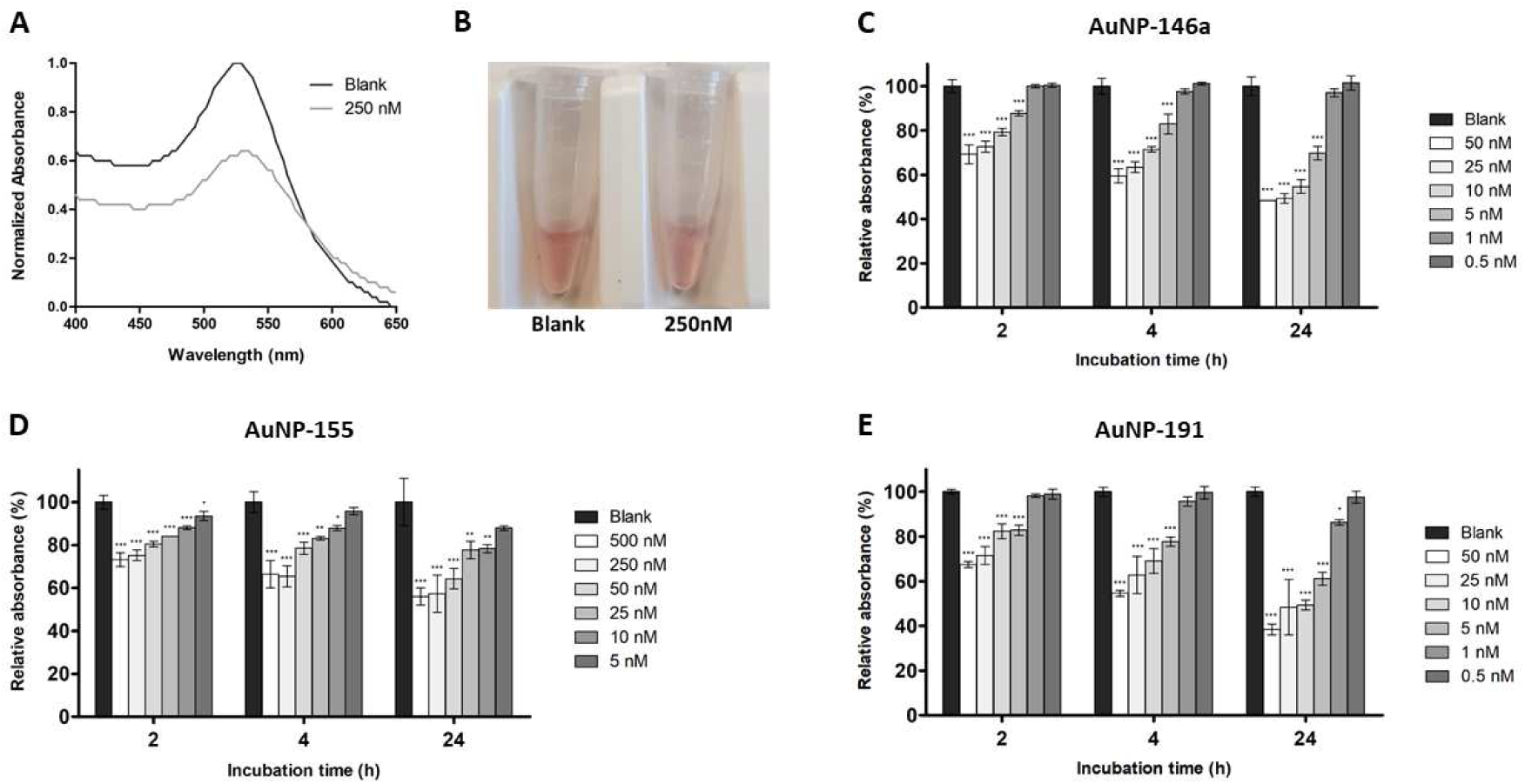
(**A**) Absorbance spectra of the sensors after 3h incubation with 250 nM of target RNA sequence (light grey), compared to the blank (no RNA) solution (dark grey). (**B**) Photograph of the samples represented in figure **A**. Data are shown for AuNP-155, representative of all the nanostructures, which had similar results. (**C-E**) LoD determination for (**C**) AuNP-146a, (**D**) AuNP-155, and (**E**) AuNP-191. For each miRNA sensor, plots of the UV-Vis absorbance maxima at around 530 nm are shown. The RNA sequences of miR-146a, 155, and 191 were added at the indicated concentrations. Presented data were collected after 2 h, 4 h, and 24 h incubation with the target (most relevant time points). Results are shown as Mean ± SD (n=3) and relative (in percentage) to the blank value (no RNA). (·) denotes a significant difference when p < 0.1, compared to the value of the blank samples; (*) when p < 0.05; (**) when p < 0.01; (***) when p < 0.001. Statistical analyses were performed by one-way ANOVA with Tukey’s *post hoc* test.

Then, the sensitivity of the three detection systems was evaluated, assessing the three sensors (AuNP-146a, AuNP-155, and AuNP-191) in the presence of their corresponding target miRNA sequence (miR-146a, miR-155, and miR-191) at different concentrations **(Figure 3C-E)**. Absorbance measurements were performed after 1, 2, 3, 4, 6, 8, and 24 h incubation with the targets. After 2 h of incubation (without any further intervention apart from absorbance measurements), the three systems were able to detect their complementary sequences at least at 10 nM (1.6 pmoles of target molecules), highlighting the versatility of the approach. Remarkably, AuNP-146a and AuNP-191 detected their targets at 5 nM (0.8 pmoles), while miR-191 was also detectable at 1 nM (160 fmoles) after 24 h. The precipitation of the AuNPs could also be seen with the naked eye down to 5 nM concentrations (not shown). The limit of detection (LoD) obtained for these sensors was similar to that observed in previous studies, in the low nanomolar range.^33,34^

In summary, the proposed sensors showed good sensitivity for the direct detection of short RNA sequences in the nanomolar range, corresponding to a few pmoles of the nucleic acids.

Since biological samples constitute a complex mixture, where many different biomolecules and other miRNAs are present, we next sought to evaluate the selectivity of the proposed sensors in the presence of multiple RNA sequences. Three RNA scramble sequences (S-1 to S-3 in **Table 1**) were added to the reaction mixture along with the respective target, corresponding to sequences of other miRNAs and siRNAs. In the case of AuNP-146a and AuNP-191, the selectivity was evaluated at 50 and 5 nM of each miRNA, while for AuNP-155 the selectivity was assessed at 500 and 10 nM. The results (Supporting Information, **Figure S2**) revealed that all three systems detected their corresponding target sequence with high selectivity across different concentrations, even when mixed with other sequences. Scramble sequences alone produced signals that were similar to the blank samples, confirming the high selectivity of the miRNA sensors.

#### 3.1.3. Simultaneous detection of three miRNAs: multiplexing

The results shown above demonstrated that the developed systems present good sensitivity and selectivity, two critical parameters in the design of miRNA sensors, due to their low abundance and sequence similarity characteristics. Nevertheless, the signature of several diseases or activation states involves the dysregulation of more than one miRNA at a time, and for that reason, multiplexing systems are desired to ease diagnosis.^20^ However, most AuNP-based sensors reported in the literature have focused on the detection of single miRNAs.^22^

The potential of this approach to detect multiple miRNAs was therefore studied. In this case, AuNPs were modified with the three oligonucleotide probes (MB-146a, MB-155, and MB-191), each one constituting 1/3 of the total MB quantity added (AuNP-Multi). This “hybrid” sensor was used to detect a mixture of the three target miRNAs (miR-146a, miR-155, and miR-191), where each one constituted 1/3 of the total quantity of target sequences. Each one of the target sequences was added at the following concentrations: 3.3, 1.7, 0.3 and 0.17 nM; making up respectively 10, 5, 1 and 0.5 nM of final concentration of targets in the sample. The results are shown in **Figure 4**.

**Figure 4.**
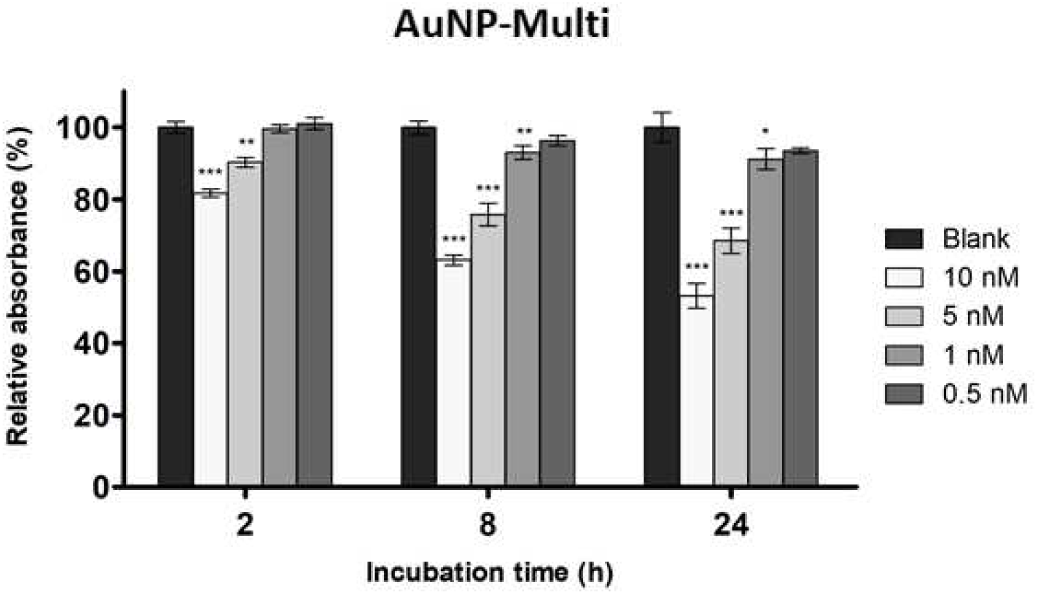
Multiplexed detection of the targets miR-146a, miR-155, and miR-191, using AuNP-Multi. Plots of the UV-Vis absorbance maxima at around 530 nm are shown. The indicated concentrations reflect the final concentration of targets in each sample, where each miRNA constitutes 1/3 of the indicated concentration. Presented data were collected after 2 h, 8 h, and 24 h incubation with the targets (most relevant time points). Results are shown as Mean ± SD (n=3) and relative (in percentage) to the blank value (no RNA). (*) denotes a significant difference when p < 0.05, compared to the value of the blank samples; (**) when p < 0.01; (***) when p < 0.001. Statistical analyses were performed by one-way ANOVA with Tukey’s *post hoc* test.

Similar to the previous individual target detection experiments, UV-Vis spectroscopy monitoring showed a decrease in absorbance maxima with increasing concentration of the target oligonucleotides (**Figure 4**). The aggregation of the AuNPs was observed after 2 h for the 10 and 5 nM samples. Longer incubation made the aggregation more evident, which also allowed the detection of the samples at 1 nM after 8 h. Remarkably, the LoD of AuNP-Multi improved with respect to the individual systems, as it was observed in previous studies.^34^

This tendency was also observed in the detection of 10, 5, 1, and 0.5 nM concentrations of the individual targets miR-146a, miR-155, and miR-191, using AuNP-Multi (**Figure S3**). The system could detect the three miRNA sequences, and the LoDs were improved, compared to the use of single oligonucleotide probe nanoparticles, reaching values of 0.5-1 nM. This improvement in the sensitivity of the AuNP-Multi system, compared to the individual sensors, could be due to the unfolding of oligonucleotide probes (MBs) on many different nanoparticles instead of unfolding many strands in less number of nanoparticles, which led to an increase of the hydrophobic interactions between the particles and the corresponding aggregation.

Due to the improved performance of the AuNP-Multi system both in the presence of single and multiple miRNAs, it was difficult to infer contributions of each MB in multiplexed AuNP-Multi-based detection. If close to equal, taking into account that each individual target is at 1/3 of the total concentration of targets, it is possible to conclude that the sensitivity of AuNP-Multi for each individual miRNA has been significantly improved (three-fold), reaching a LoD of around 300 pM (53 fmoles) after 8 h.

So, adding to its sensitivity and selectivity, the system was suitable for multiplexed detection at picomolar concentrations, corresponding to fmol levels of target sequences. This gain in sensitivity is very relevant, given the low concentrations of miRNAs in biological samples. Moreover, the sensor could be prepared to detect additional sequences simultaneously, which might increase the sensitivity even further.

The sensitive discrimination of the behavior of different biomarkers at a time, through multiplexing, is desired for miRNA-based diagnosis, since it requires the variation of several miRNAs.^20^ Also, the detection of multiple targets from the same sample decreases the variability due to sample manipulation and/or storage. With this system, several miRNAs can be detected simultaneously in the same sample, individually or in a mixture, using a single reagent, and at low concentrations. This approach would allow for a faster and more reliable identification of diseased patients, helping in diagnosis and stratification.

#### 3.1.4. Multiplexed detection: stability, detection speed, and selectivity

The activity of the AuNP-Multi sensor was studied overtime, to ensure its stability when stored at 4 °C. The same batch of AuNP-Multi was incubated for 2 h with 250 nM of the targets miR-146a, miR-155 and miR-191, i.e., 83.3 nM of each target, on different days throughout a storage time of 90 days, followed by absorbance measurements (**Figure S4**).The absorbance showed a reduction to around 80% of the blank, and AuNP aggregation could be observed clearly (not shown). Thus, the system’s performance was not affected for at least three months, highlighting its robustness, which is essential for its possible clinical use. In previous studies, storage of the sensors at RT for periods longer than 15 days affected their subsequent performance, unlike storage at 4 °C, being advisable to store the sensors at this temperature.^34^

To reduce detection time, the performance of the multiplexed sensor was studied after a short incubation with the targets, followed by low-speed centrifugation. This process could accelerate the sedimentation of aggregated AuNPs compared to the previous set-up (only gravity-dependent). The incubation time was reduced to 30 minutes, after which the samples were centrifuged at 5000 rpm for 5 minutes, and the absorbance of the supernatant was measured. The sensor was incubated in the presence of 100, 10, 5, and 1 nM of target miRNAs, meaning that each sequence was added at the concentrations of 33.3, 3.3, 1.7 and 0.3 nM. It was possible to obtain faster and reliable results, detecting at least 5 nM of miRNAs (**Figure S5**), with a significant reduction of the assay time that could facilitate its clinical implementation.

Then, the feasibility of the multiplexed detection system (AuNP-Multi) was also examined in the presence of six scramble sequences (S-1 to S-6 in **Table 1**). Similarly to selectivity assays, the experiments included blank samples, samples containing the three targets mixed with the six scramble sequences, only the targets, or only the scramble sequences (**Figure S6**). The studies were performed at the concentrations of 10, 5 and 1 nM, so each target was added at the concentrations of 3.3, 1.7, and 0.3 nM. For scramble sequences, in turn, each one was added at the same concentration of the targets, thus making up respectively 20, 10 and 2 nM final concentration of scramble sequences. In this case, the aggregation of the AuNPs also occurred selectively, and only the presence of the target sequences for the concentrations of 10 nM and 5 nM, as shown by the UV-Vis absorbance measurements (**Figure S6**). In this case, no aggregation was observed for the 1 nM concentration, due the low differences affecting sensitivity and selectivity studies at the proximity of the LoD.^33^ In addition, sensitivity and selectivity at 1 nM had been achieved with this system when using DNA sequences as targets (data not shown).

In general, the AuNP-Multi system was selective for the presence of the three target miRNA sequences, discriminating them from a mixture of diverse sequences at concentrations as low as 1-5 nM. These results further validated the system for multiplexed detection of miRNAs in complex samples and its possible application in miRNA profiling.

Since the sensors presented herein are based on color changes, they could be a promising PoC system, due to their simple readout, without the need for sophisticated or expensive equipment, allowing for simple, fast, and cost-effective detection.^63^ However, the absorbance differences are reduced when detecting samples at low concentrations, still requiring the use of spectrophotometers or absorbance readers to detect those differences. Such equipment might not be readily available in PoC settings, in particular, those with limited resources. LFAs could overcome these limitations, offering advantages like improved detection speed, portability, low cost, and ease of use. In addition, LFAs can be more easily integrated with smartphone-based readers, facilitating real-time data interpretation and analysis.^64,65^ Thus, the proposed system could benefit substantially from adapting to the LFA format, improving its applicability for PoC detection.

On the other hand, although these sensors could detect specific miRNAs at picomolar concentrations, miRNA concentrations in biological samples are several orders of magnitude lower. For instance, in serum/plasma samples, individual miRNAs can range from 25 to 8000 copies per µ L, corresponding to concentrations between 40 aM and 15 fM.^20,66^ In addition, the incubation of the AuNPs with the target sequences or biological samples would require further dilutions, to include the assay buffer and the AuNPs. Thus, direct detection with this system, although in multiplexed forms, would be insufficient to detect miRNAs in clinical samples.

In this regard, nucleic acid amplification techniques have been sought to increase the sensitivity of AuNP-based detection strategies.^22^ Since the RT-PCR gold standard is not straightforward for the amplification of miRNAs, due to their short length, a wide range of alternative isothermal amplification methods have been proposed for miRNA detection. These techniques can amplify short nucleic acids with efficiencies comparable to PCR, but occurring at a single temperature, avoiding the need for thermal cycling. Indeed, these reactions only require a thermal block (or even water bath), offering an additional advantage for PoC detection.^67,68^ In this regard, coupling to isothermal amplification could be explored to improve the sensitivity of the system, allowing PoC detection in relevant samples. LFA and isothermal amplification are explored in the following sections to improve the proposed system.

### 3.2. AuNP-based lateral flow sensors

3.2.1. **Preparation and characterization of the sensors**

In order to adapt the system to a LFA format, more suitable for PoC detection, terminal cholesterol moieties were replaced with biotin ones, taking advantage of its high affinity to the streptavidin protein, which can be deposited in lateral flow strips (**Figure 1C**).

For the preparation of the sensors, NC strips were prepared with a streptavidin test line. The protocol for deposition of the streptavidin protein was optimized for maximum signal-to-noise ratio, as well as the assay buffer, both described in *Section 2* (data not shown). In parallel, commercial citrate-stabilized AuNPs of 40 nm were modified with hairpin oligonucleotides that contained the thiol moiety at one end and biotin at the other, optimizing the functionalization conditions. The resulting AuNPs were characterized as described for sensors in solution (**Figure S7**).

TEM images showed a higher morphological diversity for 40 nm AuNPs, with a broader size distribution and a mean diameter of 38.1 ± 5.9 nm (**Figure S7A** and **B**). The AuNPs were modified with the oligonucleotides MB-146a*, MB-155*, and MB-191* (**Table 1**), forming AuNP-146a, AuNP-155, AuNP-191, and AuNP-Multi, as described above. In this case, the loading of the AuNPs was reduced by avoiding the addition of NaCl, while approximately 5% of the AuNP surface was covered with LP-PEG, used to stabilize the nanostructure. It was observed in initial experiments (data not shown) that a high MB loading on the surface of AuNPs produced unspecific signals on test strips, due to non-specific opening of the MBs and exposure of the hydrophobic biotin molecule, consistent with previous reports.^36,37^ The loading efficiency was reduced to around 25% (**Figure S7C**), in agreement with the method used.^69^

DLS, ELS, and UV-Vis spectroscopy results were also consistent with the desired functionalization. The hydrodynamic diameter increased from 46 nm to 50-55 nm, with a PDI of 0.2 (**Figure S7D**); the Zeta potential increased from -40 mV to -20 mV (**Figure S7E**); and the absorbance spectra were similar (**Figure S7F**). All these results pointed to the correct functionalization of the AuNPs with biotin-containing MBs. The differences in the results when compared to cholesterol-containing counterparts indicated the lower MB loading on the AuNP surface attained in this case.

#### 3.2.2. Sensor performance: sensitivity and selectivity

LFA sensors are based on the exposure of the biotin moiety, due to the opening of the MB structure upon binding of the target. Then, loading of the mixture onto streptavidin lateral flow strips leads to the retention of the AuNPs on the test line, producing a colored signal.^36,37^

The LoD of the prepared sensing systems (AuNP-146a, AuNP-155, and AuNP-191) was studied by incubating with different concentrations of the target miRNAs (miR-146a, miR-155, and miR-191), loading the mixtures onto the test strips, and quantifying the colored signal. First, a broad range of target concentrations (from 500 nM to 1 nM) was evaluated (**Figure 5A-C**), showing that all the miRNA sequences were detected at least at 10 nM (200 fmoles of target molecules). Then, a range of concentrations between 10 nM and 1 nM was tested for precise determination of the LoD under 10 nM (**Figure 5D-F**). Notably, AuNP-146a and AuNP-191 detected their targets at 5 nM (100 fmoles), while miR-191 was also detectable at 2.5 nM (50 fmoles) after 24 h. Previously reported systems had LoDs of around 0.5 nM.^36,37^ Therefore, LFA sensors could also detect their targets in the nanomolar range, which in this case corresponded to fmol quantities of nucleic acids.

**Figure 5.**
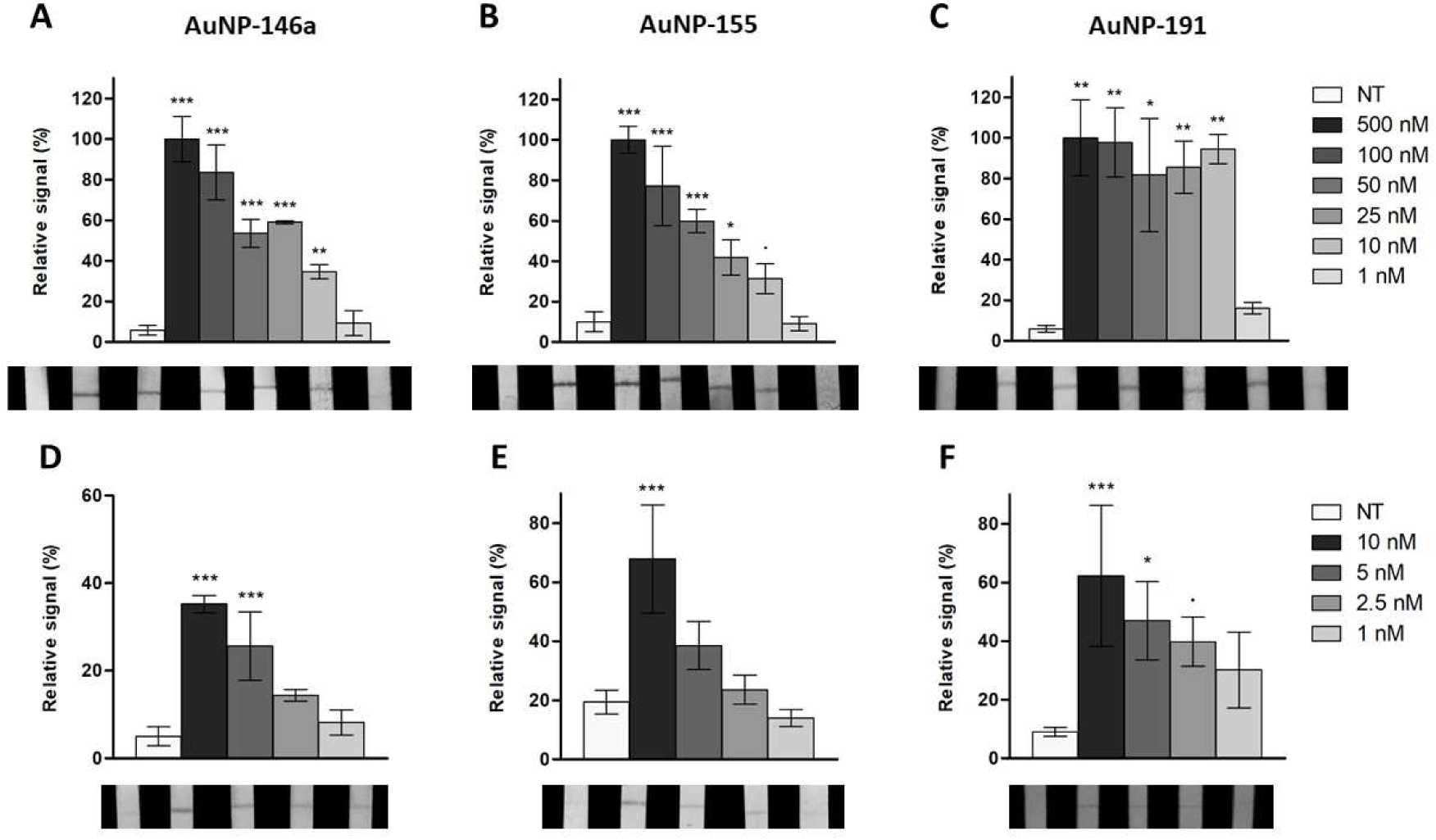
LoD determination of (**A,D**) AuNP-146a, (**B,E**) AuNP-155, and (**C,F**) AuNP-191, using streptavidin NC strips – photographs of the strips and signal intensity plots. From left to right were added: a negative control (NT – no target), and the corresponding target sequence at different concentrations. (**A,B,C**) Concentrations from 500 nM to 1 nM (see legend). (**D,E,F**) Concentrations between 10 nM and 1 nM (see legend). The plots show the integrated signal for each NC strip, as Mean ± SD (n=3) and relative (in percentage) to the highest intensity value (500 nM). (·) denotes a significant difference when p < 0.1, compared to the negative control (NT); (*) when p < 0.05; (**) when p < 0.01; (***) when p < 0.001. Statistical analyses were performed by one-way ANOVA with Tukey’s *post hoc* test.

The evaluation of these sensors continued with selectivity studies in the presence of multiple RNA sequences. The RNA scramble sequences S-1 to S-3 (**Table 1**) were added to the reaction mixtures along with the targets, at concentrations of 500 nM and 10 nM (**Figure S8**). The results showed that the three systems were able to detect their targets with high selectivity, producing significant colored signals only when the corresponding target was present, either alone or mixed with other sequences, which did not produce signals by themselves. These data confirmed the good selectivity presented as well by LFA sensors.

#### 3.2.3. Simultaneous detection of three miRNAs: multiplexing

Since the proposed sensors presented good sensitivity and selectivity, which are important for miRNA detection, the ability of this approach for multiplexed detection was studied next. The biotin-containing version of the AuNP-Multi system was prepared and challenged with an equal mixture of the three target miRNAs. For the first broad range of concentrations, each one of the target sequences was added at 167, 33, 17, 8.3, 3.3, and 0.3 nM; making up respectively 500, 100, 50, 25, 10, and 1 nM of the final concentration of targets in the sample. Then, each target was added at 3.3, 1.7, 0.8, and 0.3 nM; making up 10, 5, 2.5, and 1 nM of final concentrations, respectively, for precise LoD determination. The results are shown in **Figure 6**.

**Figure 6.**
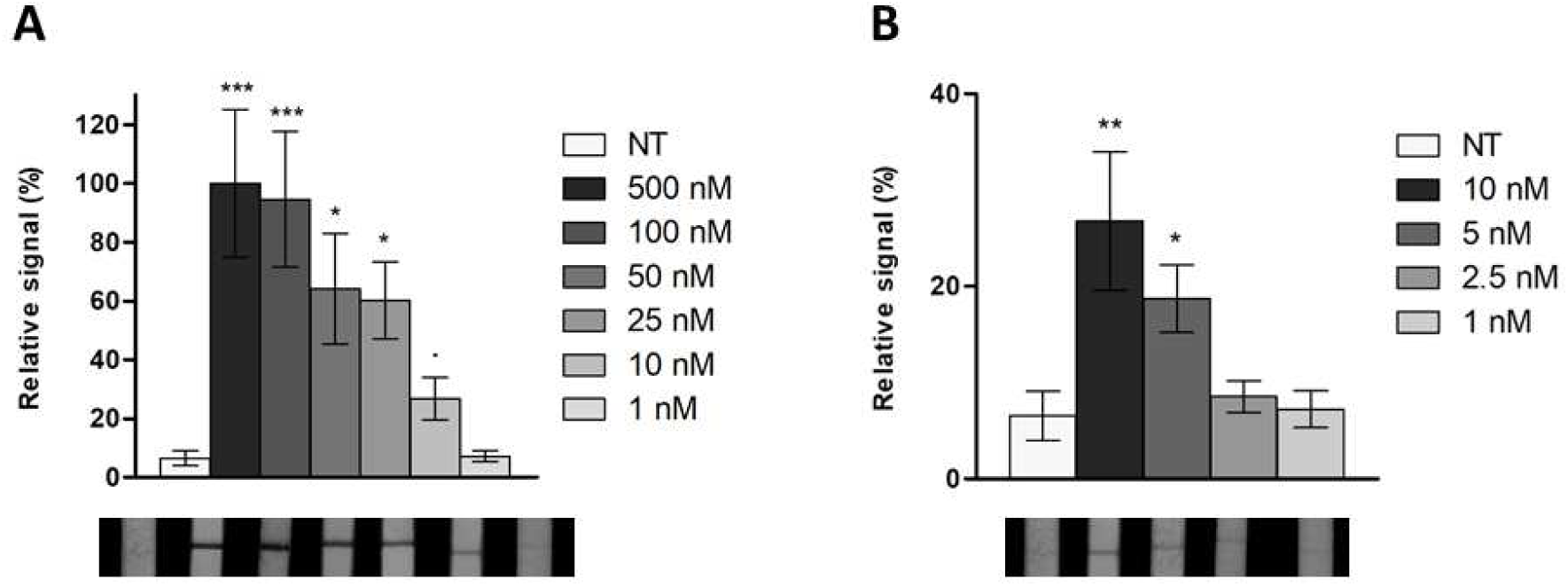
Multiplexed detection of the targets miR-146a, miR-155, and miR-191, using AuNP-Multi and streptavidin NC strips – photographs of the strips and signal intensity plots. From left to right were added: a negative control (NT – no target); and the three target sequences at different concentrations. The indicated concentrations reflect the final concentration of targets in each sample, where each miRNA constitutes 1/3 of the indicated concentration. (**A**) Concentrations from 500 nM to 1 nM (see legend). (**B**) Concentrations between 10 nM and 1 nM (see legend). The plots show the integrated signal for each NC strip, as Mean ± SD (n=3) and relative (in percentage) to the highest intensity value (500 nM). (·) denotes a significant difference when p < 0.1, compared to the negative control (NT); (*) when p < 0.05; (**) when p < 0.01; (***) when p < 0.001. Statistical analyses were performed by one-way ANOVA with Tukey’s *post hoc* test.

The AuNP-Multi system had a LoD between 10 nM and 1 nM (**Figure 6A**), like the biotin-containing individual sensors. The study in the range of 10 nM to 1 nM revealed that the LoD was around 5 nM, an intermediate value to those obtained for the individual sensors (**Figure 6B**). These results hinted towards a possible almost equal contribution of all the MBs for the detection of the three target sequences, meaning that the system reached a LoD of around 1.7 nM (30 fmol) for each miRNA.

In this case, sensitivity improvements were not observed when using the biotin-containing version of AuNP-Multi and LFA detection, apart from the effect of using 1/3 of the concentration of each miRNA. Unlike the system in solution, LFA sensors depend on the number of biotin moieties available to interact with streptavidin on the NC strips, with a 1:1 interaction ratio with each binding site on the streptavidin molecules. In turn, the system in solution is based on the aggregation of the AuNPs between each other, and depends on the number of AuNPs that are exposing the cholesterol moieties. Thus, an increase in this number would promote AuNP aggregation, but could have a negligible effect on LFAs.

Still, the LFA sensor retained the applications and advantages of multiplexed systems, discussed above, allowing the detection of several miRNAs simultaneously in the same sample, at nanomolar concentrations, corresponding to fmol quantities.

A possible future development of this approach could be a multiplexed lateral flow format, though the modification of each MB with a different reporter molecule (e.g., biotin, digoxigenin, fluorescein), and placement of different biorecognition molecules (e.g., streptavidin, antibodies, aptamers) in separate test lines of a lateral flow sensor. With this, individual identification of each miRNA biomarker, along with individual quantification, could be possible in a single assay, but assessing the dysregulation of multiple miRNAs.^70,71^

Finally, the performance of AuNP-Multi was evaluated in the presence of the six scramble sequences (S-1 to S-6 in **Table 1**). Like previously, negative controls, samples containing the three targets mixed with the six scramble sequences, only the targets, or only the scramble sequences were prepared. The studies were performed at 500 and 10 nM, so each target was added at 167 and 3.3 nM, respectively. For scramble sequences, in turn, each one was added at the same concentration of each target, thus making up respectively 1000 nM and 20 nM total concentration of scramble sequences. The results showed AuNP-Multi was selective for the presence of the three target miRNA sequences, either alone or in the presence of scramble sequences, being able to discriminate them from the scramble mixture (**Figure S9**). Thus, the AuNP-Multi system was also suitable for selective multiplexed detection at low nanomolar concentrations in the LFA format.

The results obtained for the LFA sensors showed these systems could reproduce the merits showed previously for colorimetric sensors (in solution), being a promising system for miRNA-based diagnosis and molecular profiling. Notably, the use of a LFA format improves significantly the applicability of the sensors in PoC detection, especially in resource-limited facilities. However, its nanomolar LoDs are insufficient to detect miRNAs in biological samples. Isothermal amplification techniques have been coupled to LFAs for miRNA detection, retaining the advantages of PoC devices.^22,72^ Based on these reports, one of these approaches, the Exponential Amplification Reaction (EXPAR), was explored to improve the sensitivity of LFA sensors for detection in relevant samples.

### 3.3. EXPAR amplification and detection of miRNAs

Exponential Amplification Reaction (EXPAR) was the isothermal method used in this work for signal amplification, coupled to LFA detection. Short oligonucleotides, like miRNAs, can be amplified 10^6^-10^9^ fold by EXPAR, in a few minutes, through an isothermal reaction, detecting as low as 10 copies of miRNA per µ L (low aM concentrations). This technique consists of the use of a DNA template that can trigger successive steps of miRNA extension, nicking, and release of an analog sequence, which is continuously amplified (**Figure 7**). Like qPCR, the EXPAR reaction can be monitored in real-time using a SYBR Green dye, which binds to the target-template duplexes and double-stranded sequences extended from them. Nevertheless, unlike qPCR, the EXPAR product continues to be generated when the fluorescence plateau is reached.^38–42^

**Figure 7.**
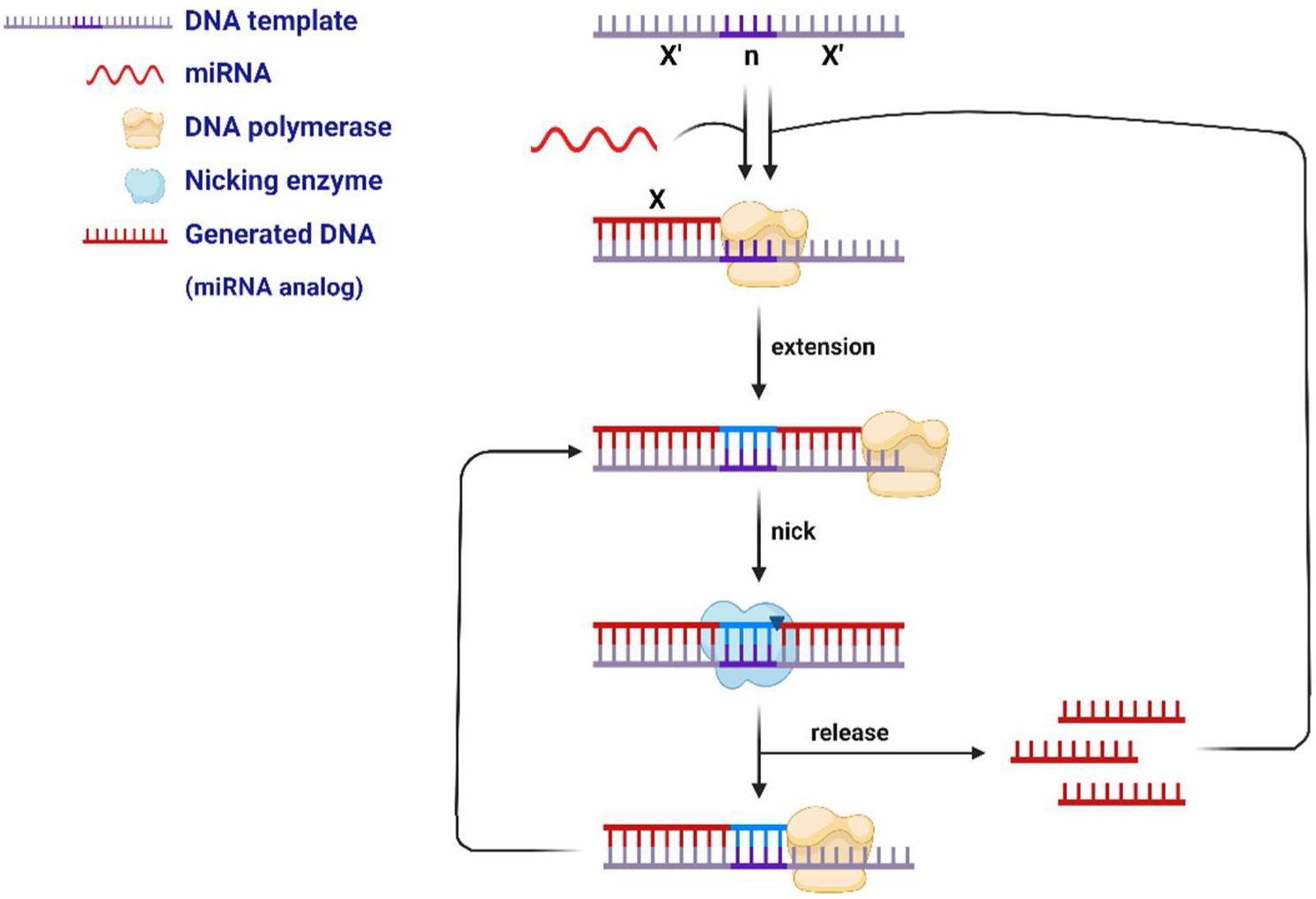
Schematic representation of the EXPAR amplification. The system is composed of a DNA template, which contains two regions (X’) that are complementary to a part of the target miRNA (X), separated by a region that can be cut by a nicking endonuclease (n). The miRNA binds to the DNA template, and a polymerase extends the sequence, generating the double-stranded recognition site of the nicking endonuclease. This enzyme cuts the extended strand, releasing a sequence that is analog to the miRNA (and shorter), which reinitiates the process. This sequence is generated continuously, producing an exponential amplification of the initial target sequence. Image created with BioRender.com

Different template configurations can be used in the EXPAR reaction,^73^ while the form X’nX’ (**Figure 7**) is the most studied and was used in this work to detect the miRNA 155. X’nX’ templates are typically composed of two 10-20 nt sequences (X’) complementary to the target (X) to be detected, and separated by the recognition site of a nicking endonuclease (n). In this case, the nicking enzyme Nt.BstNBI was used, which recognizes the double-stranded 5’-CAGTC-3’ sequence, generated upon miRNA/analog extension, and nicks the strand 4 nt upstream of the recognition sequence. This 4 nt “spacer” was also included in the template sequence (Temp, **Table 2**). A DNA-based analog of miR-155 was used as the target X sequence (T-155, **Table 2**).

The design of the template and the reaction protocol were optimized before coupling with the LFA system, by monitoring the reaction in a real-time PCR thermal cycler. EXPAR reactions are particularly affected by non-specific background amplification, and modifications to the original protocol have been proposed to minimize its effect. These variations help achieve sufficient time separation between the appearance of specific and non-specific amplification signals.^39,73,74^ The optimization process (not shown) included the adjustment of parameters like: the length of the X’ region of the template, which was shortened to 12 nucleotides; capping of the 3’ end of the template with phosphate groups; physical separation of the template from the polymerase until reaction start; lowering of the reaction temperature from 55 °C to 51 °C; and pre-incubating the reaction parts at 51 °C before mixing them (hot-start reaction). The final reaction protocol is described in *Section 2*. Remarkably, under optimized conditions, fluorescence monitoring of the EXPAR reaction showed that at least 100 fM of the target T-155 could be detected (**Figure 8A**). In addition, preliminary detection experiments at 50 °C (data not shown) revealed that amplification down to 1 fM could be possible. Also, ∼10 aM miRNA concentrations have been detected in previous reports.^40^

**Figure 8.**
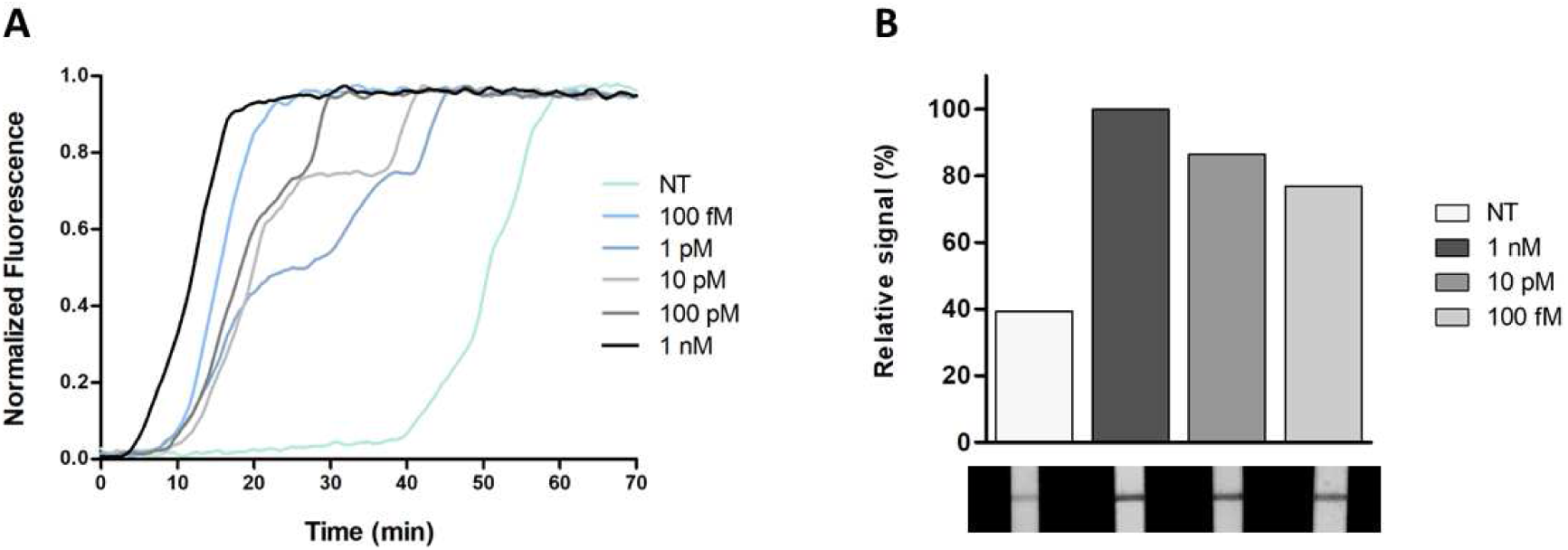
(**A**) Fluorescence curves obtained from real-time monitoring of the EXPAR reaction – variation of SYBR Green fluorescence over time (normalized signal). The T-155 sequence was added at concentrations of 1 nM, 100 pM, 10 pM, 1 pM, and 100 fM, compared with samples without the target sequence (NT – no target) (see legend). The plots represent the mean of 4 replicas. (**B**) Detection of the T-EXP sequence after a 15 minutes EXPAR reaction, using AuNP-EXP and streptavidin NC strips – photographs of the strips and signal intensity plots. The T-155 sequence was added at the concentrations of 1 nM, 10 pM, and 100 fM, compared to samples without target sequence (NT – no target). The plots show the integrated signal for each NC strip and relative (in percentage) to the highest intensity value (1 nM).

For coupling to LFA detection, 40 nm AuNPs were modified to detect the EXPAR product, using the oligonucleotide MB-EXP (**Table 1**), a MB that was complementary to the EXPAR product, i.e., the generated X sequence. In this case, the amplification product consisted in a 12 nt partial sequence of T-155, designated T-EXP (**Table 1**). The resulting AuNPs were designated AuNP-EXP, and the sensitivity was studied using a synthetic T-EXP DNA sequence (**Figure S10**). The results showed that the T-EXP sequence could be detected at 10 nM. Studies at concentrations between 10 nM and 1 nM showed no change in the LoD of this system (data not shown). Nonetheless, with the exponential amplification provided by EXPAR, the final concentrations of the reaction products to be detected by lateral flow would be much higher than 10 nM, validating the system for coupling with LFA.

After this, LFA detection studies were performed to determine the optimal detection conditions, by incubating a portion of the reaction volume with AuNP-EXP after amplification and loading onto NC strips. A reaction volume of 1 µ L was incubated with the AuNPs to minimize the background signals, and showed the most significant difference between the negative and the positive samples (**Figure S11**). Although the reaction components presented some interference, it was possible to distinguish negative and positive signals without purification of the amplification products, simplifying the procedure for PoC detection. On the other hand, since the EXPAR amplification product is generated continuously and exponentially once the reaction starts, different reaction times were studied to further optimize detection conditions. The results showed that a reaction time of 15 minutes was adequate to detect all target concentrations, while distinguishing them from the negative samples (**Figure S12**). Longer reactions (30 min and above) showed signal saturation in the strips, including negative samples, indicating that shorter reaction times are necessary to avoid non-specific signals. Finally, the LFA detection study of the products of the 15-minutes EXPAR reaction revealed the samples could be detected at all the concentrations detected in fluorescence monitoring, i.e., at least at 100 fM (**Figure 8B**). Additionally, the signal seemed to be proportional to the logarithm of the initial target concentration, consistent with exponential amplification. Given the observed signal intensities, even for the 100 fM sample, the proposed LFA could easily detect lower concentrations. Further experiments will explore this possibility, but it is clear that the system seems promising for detecting miRNAs at low concentrations. EXPAR templates can be designed for the amplification and detection of multiple sequences, while LFA systems can be used in multiplexed formats, enabling miRNA profiling with PoC systems.

## 4. Conclusion

In this paper, we report the development of a sensor based on gold nanoparticles (AuNPs) and hairpin-shaped oligonucleotides, for the detection of nucleic acids in physiological samples. The recognition of target sequences alters the colloidal stability of AuNPs, producing color changes that are easily detected with the naked eye. The sensor demonstrates good sensitivity and selectivity in the detection of single miRNAs, and can also recognize specifically several miRNAs at a time. These multiplexing capabilities allow for improved sensitivity, reaching picomolar levels of nucleic acids. This system was further adapted to a lateral flow-based format, producing colored signals in nitrocellulose strips in the presence of the targets. It retained its exceptional properties, while making it more suitable for application as a PoC device. Finally, the lateral flow system was successfully coupled with EXPAR isothermal amplification, achieving sensitivities close to those required for detection in clinically relevant samples. Besides miRNA profiling, our systems can be easily tuned for the detection in liquid biopsies of any nucleic acid biomarker, extending their versatility for multiplexed nucleic acid detection in different disease situations or physiological contexts.

In summary, the sensors developed herein, coupled with isothermal amplification, hold promise for future clinical applications. It would be a fast, versatile, reliable, and cost-effective system, suitable for use in PoC settings and/or resource-limited facilities. The complete detection assay could be performed in 30 minutes, including a 15-minutes EXPAR reaction, followed by the lateral flow assay. Such fast, reliable, and affordable diagnostic systems are essential for implementing detection strategies in healthcare systems. In addition, they can ease the detection of overexpressed nucleic acid biomarkers related to physiological processes. Finally, the possibility of performing rapid liquid biopsy analyses in the PoC could largely contribute to the fast and early detection of diseased patients, and to better prognostic follow-up, increasing the chances of successful treatment and survival.

## Acknowledgements

This work was supported by the Spanish Ministry of Science, Innovation and Universities (SAF2017-87305-R, PID2020-119352RB-I00, PID2023-146982OB-I00), ‘Severo Ochoa’ Programme for Centres of Excellence in R&D (CEX2020-001039-S), Comunidad de Madrid (REACT-NANOCOV, S2022/BMD-7403 RENIM-CM), and Asociación Española Contra el Cáncer. Catarina Coutinho acknowledges the financial support from Asociación Española Contra el Cáncer (PRDMA18030CAST).

## Supporting Information

Supporting Information is available from the author.

## Conflicts of interest

There are no conflicts to declare.

## Notes

### Competing Interest Statement

The authors have declared no competing interest.

