## Supporting Information for "Hairpin-Functionalized Gold Nanoparticles as an Adaptable Platform for Detecting MicroRNA Signatures"

### 1. Synthesis of lipoic acid-modified polyethylene glycol (LP-PEG)

Lipoic acid-modified polyethylene glycol (LP-PEG) (compound **2**) was synthesized by the activation of lipoic acid (ABCR GmbH, Karlsruhe, Germany) to form compound **1**, followed by amide bond formation (**Figure S1**).[1,2] The reagents *N*-hydroxysucciminide (NHS) and *N,N'*-dicyclohexylcarbodiimide (DCC) were obtained from Sigma Aldrich (San Luis, MO, USA), while $H_2N$-PEG-OH of 3 KDa was obtained from Iris Biotech (Marktredwitz, Germany). The solvents tetrahydrofuran (THF) and ethyl acetate were purchased from Scharlab (Barcelona, Spain). The compounds were analyzed by $^{1}H$ and $^{13}C$ nuclear magnetic resonance (NMR), using a Bruker DPX (400 MHz) spectrometer (Bruker, Mannheim, Germany). Mass quantification was performed at SIdI/UAM (Madrid, Spain), using electronic ionization (EI) and matrix-assisted laser ionization (MALDI).

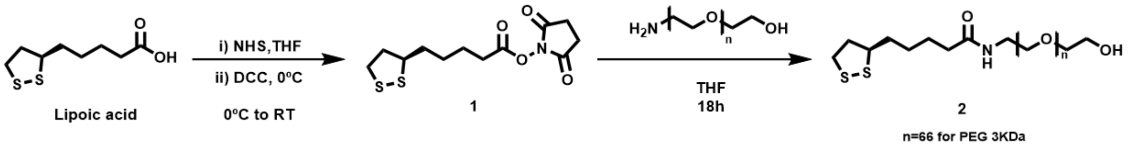

**Figure S1.** Schematic representation of the synthesis of lipoic acid-modified polyethylene glycol (LP-PEG) of 3 KDa (compound **2**).

To obtain compound **1**, lipoic acid (1 equiv.) and NHS (1.2 equiv.) were dissolved in THF (0.5 M), and the solution was stirred at 0 °C for 10 min. Then, DCC (1.2 equiv.) was dissolved in THF (3.5 M) and added slowly to the previous solution. The reaction was stirred at room temperature (RT) for 5 h. The mixture was filtered using filter paper, and the solid was washed with cold ethyl acetate. The solvent was evaporated under vacuum, obtaining the desired compound **1** as a yellow oil (yield of 98%).[1,2] $^{1}H$ NMR ($CDCl_3$, 400 MHz): δ 3.54 (dq, 1H), 3.11 (m, 2H), 2.8 (s, 4H), 2.57 (t, 2H), 2.41 (m, 1H), 1.88 (m, 1H), 1.72 (m, 2H), 1.67 (m, 2H), 1.52 (m, 2H). $^{13}C$ NMR ($CDCl_3$, 101 MHz): δ 169.27, 168.44, 56.19, 40.25, 38.73, 34.78, 30.87, 28.32, 25.69, 24.45. MS (EI): *m/z* calculated for $C_{12}H_{17}NO_4S_2$ ($M^+$) 303.05, found 303.05.[2]

Finally, to LP-PEG (compound **2**), $H_2N$-PEG-OH of 3 KDa (1 equiv.) and the lipoic derivative **1** (2 equiv.) were dissolved in THF, and the reaction was stirred for 18 h. The crude was purified by dialysis using 3.5 KDa tubing membranes (Spectrum,

ThermoFisher Scientific, Waltham, MA, USA) against distilled water. After stirring, the desired product was obtained as a yellowish solid (yield of 45%).[1,2] $^1$H NMR ($CDCl_3$, 400 MHz): δ 3.74 (s, 258H), 3.42 (t, 2H), 3.24 (dd.47 (m, 1H), 2.53 (t, 2H), 2.3 (t,2H), 2.03 (m, 1H), 1.79 (m, 1H), 1.66 (m, 4H), 1.46 (m, 2H). $^{13}$C NMR ($CDCl_3$, 400 MHz): δ 173.1, 171.9, 70.6, 56.4, 40.3, 39.28, 38.5, 36.3, 34.7, 33.51, 29.8, 25.5, 24.8. MS (MALDI): *m/z* calculated for $C_8H_{13}ONS_2(PEG)_{66}$ 3047.7, found 3046.8.[2]

### 2. AuNP-based colorimetric sensors in solution

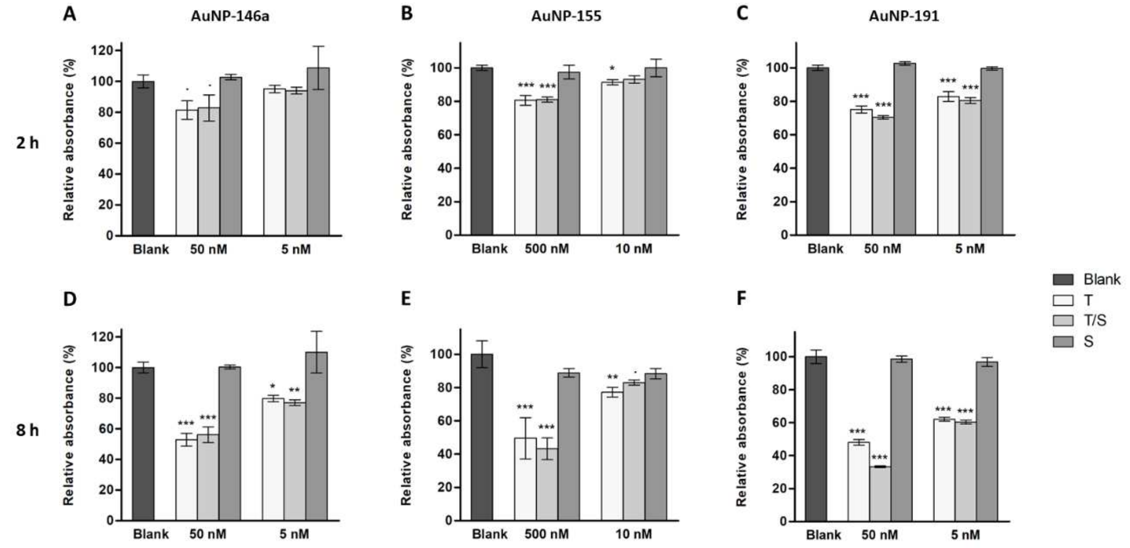

**Figure S2.** Selectivity studies of (**A,D**) AuNP-146a, (**B,E**) AuNP-155, and (**C,F**) AuNP-191. For each miRNA sensor, plots of the UV-Vis absorbance maxima at around 530 nm are shown. In addition to a non-RNA sample (Blank), for each of the indicated concentrations, three samples were prepared: a sample containing only the target at the indicated concentration (T); a mixture of the target and three scramble sequences (S-1, S-2, and S-3), each one at the indicated concentration (T/S); a sample containing only the three scramble sequences (S). Presented data were collected after (**A,B,C**) 2 h and (**D,E,F**) 8 h incubation with the RNA sequences (most relevant time points). Results are shown as Mean ± SD (n=3) and relative (in percentage) to the blank value (no RNA). (·) denotes a significant difference when $p < 0.1$, compared to the value of the blank samples; (*) when $p < 0.05$; (**) when $p < 0.01$; (***) when $p < 0.001$. Statistical analyses were performed by one-way ANOVA with Tukey's *post hoc* test.

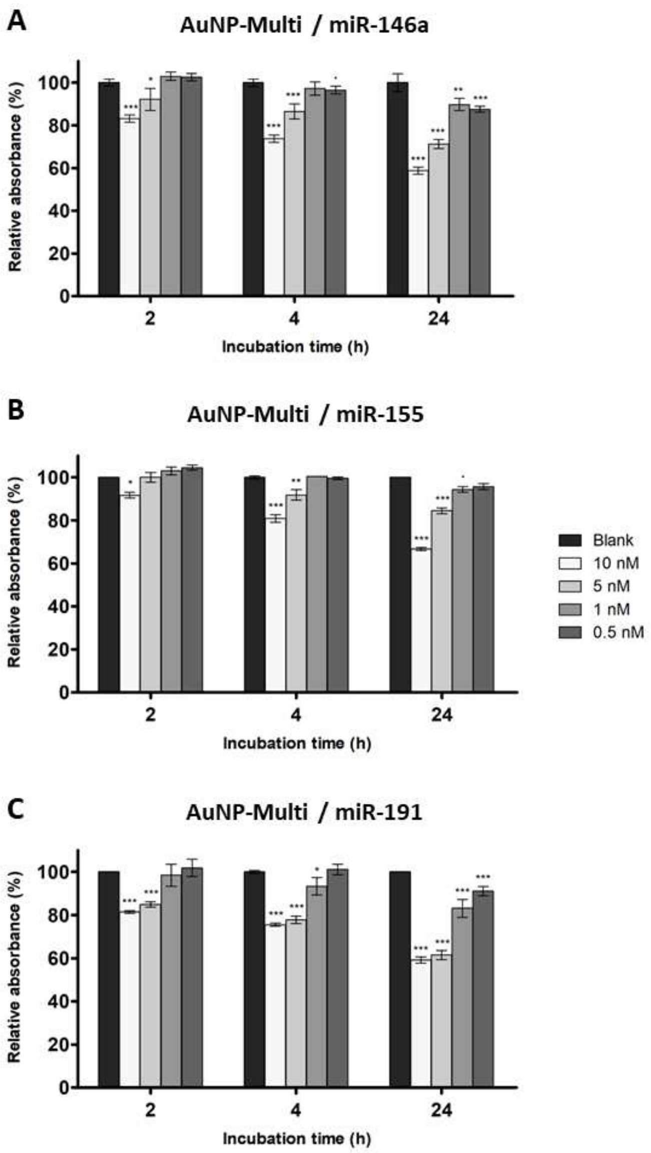

**Figure S3.** Detection of (**A**) miR-146a, (**B**) miR-155, and (**C**) miR-191, using AuNP-Multi. Plots of the UV-Vis absorbance maxima at around 530 nm are shown. The RNA sequences of miR-146a, 155, and 191 were added at the indicated concentrations. Presented data were collected after 2 h, 4 h, and 24 h incubation with the target (most relevant time points). Results are shown as Mean ± SD (n=3) and relative (in percentage) to the blank value (no RNA). (·) denotes a significant difference when $p < 0.1$, compared to the value of the blank samples; (*) when $p < 0.05$; (**) when $p < 0.01$; (***) when $p < 0.001$. Statistical analyses were performed by one-way ANOVA with Tukey's *post hoc* test.

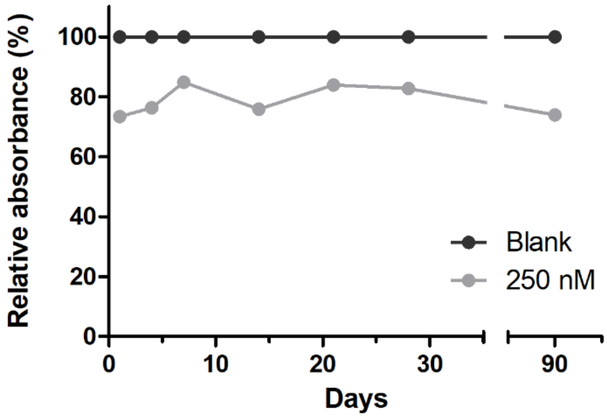

**Figure S4.** Effect of the storage time at 4 °C on the activity of the multiplexed sensor in the presence of miR-146a, miR-155, and miR-191. A batch of AuNP-Multi was incubated with 250 nM final concentration of miRNAs (83.3 nM each miRNA) on different days after storage (1, 4, 7, 14, 21, 28, and 90 days). After 2 h, the absorbance of the system was measured and the absorbance maxima at around 530 nm is shown, compared to the blank samples (no RNA). Results are presented as relative (in percentage) to the blank value.

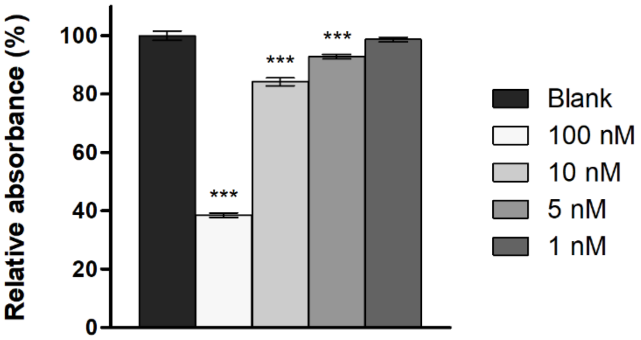

**Figure S5.** Multiplexed detection of miR-146a, miR-155, and miR-191, using AuNP-Multi after low-speed centrifugation. After 30 min incubation, the samples were centrifuged at 5000 rpm for 5 min, and the absorbance of the system was measured. UV-Vis absorbance maxima at around 530 nm are shown. The indicated concentrations reflect the final concentration of targets in each sample, where each miRNA constitutes 1/3 of the indicated concentration. Results are shown as Mean ± SD (n=3) and relative (in percentage) to the blank value (no RNA). (***) denotes a significant difference when $p < 0.001$, compared to the value of the blank samples. Statistical analyses were performed by one-way ANOVA with Tukey's *post hoc* test.

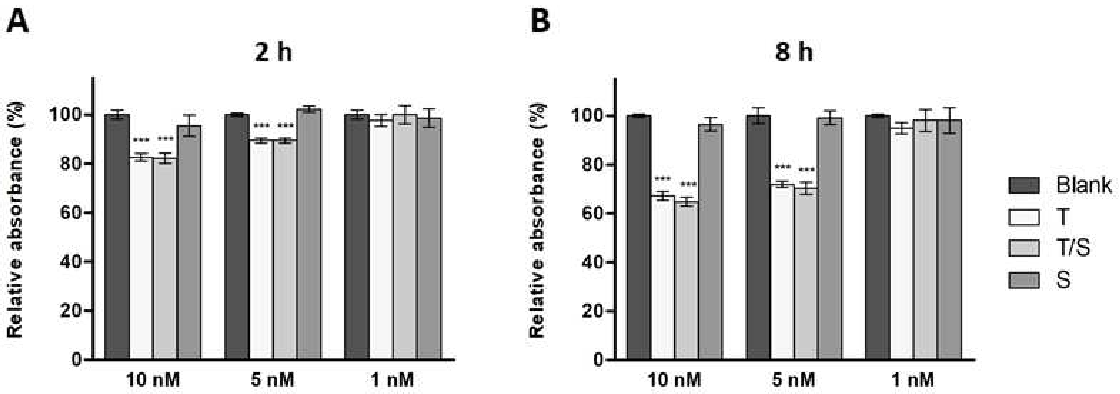

**Figure S6.** Multiplexed detection of the targets miR-146a, miR-155 and miR-191 using AuNP-Multi, in the presence of scramble sequences. Plots of the UV-Vis absorbance maxima at around 530 nm are shown. In addition to a non-RNA sample (Blank), for each of the indicated concentrations, three samples were prepared: a sample containing the three targets, at the indicated total concentration (T); a mixture of the three targets and the six scramble sequences (S-1 to S-6), each one at the same concentration of each target (T/S); a sample containing only the six scramble sequences (S). Presented data were collected after (**A**) 2 h and (**B**) 8 h incubation with the RNA sequences. Results are shown as Mean ± SD (n=3) and relative (in percentage) to the blank value. (***) denotes a significant difference when $p < 0.001$, compared to the value of the blank samples. Statistical analyses were performed by one-way ANOVA with Tukey's *post hoc* test.

### 3. AuNP-based lateral flow sensors

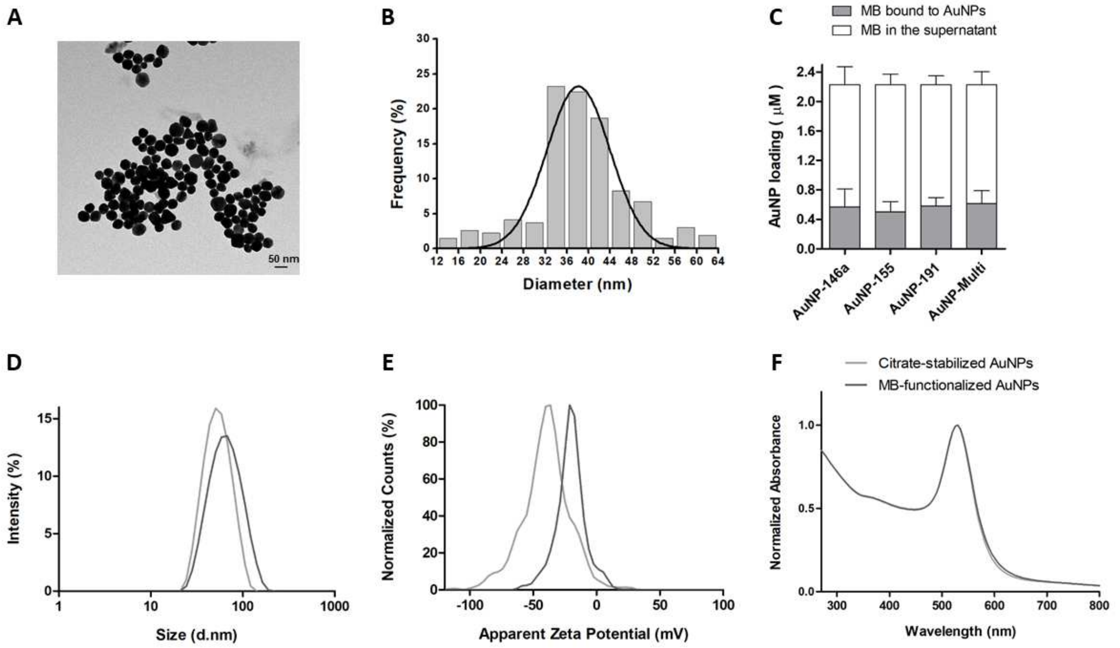

**Figure S7.** Characterization of AuNP-based lateral flow sensors. (**A**) TEM image of 40 nm citrate-stabilized AuNPs. (**B**) Size distribution obtained from TEM images, by measuring at least 200 AuNPs, using the Analyze Particles function of ImageJ software, and fitting to a Gaussian curve in Excel program. (**C**) Loading efficiency of 40 nm AuNPs with biotin-modified MBs. Results are presented as Mean ± SD: n=3 for AuNP-146a; n=4 for AuNP-155, AuNP-191, and AuNP-Multi. (**D**) Hydrodynamic diameter, (**E**) Zeta potential distribution, and (**F**) UV-Vis absorbance spectra of both citrate-stabilized (light grey) and MB-functionalized AuNPs (dark grey). Data are shown for AuNP-191, representative of all the nanostructures, which had similar features.

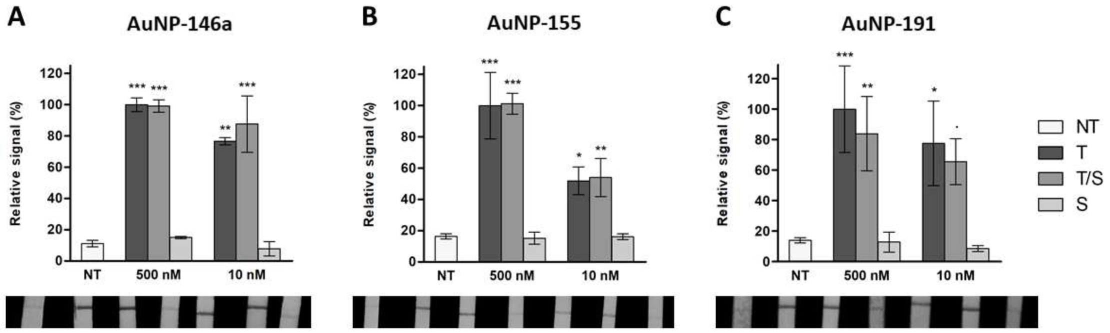

**Figure S8.** Selectivity studies of (**A**) AuNP-146a, (**B**) AuNP-155, and (**C**) AuNP-191, using streptavidin NC strips – photographs of the strips and signal intensity plots. From left to right were added: a negative control (NT – no target); a sample containing only the target at the indicated concentration (T); a mixture of the target and three scramble sequences (S-1, S-2, and S-3), each one at the indicated concentration (T/S); a sample containing only the three scramble sequences (S). The plots show the integrated signal for each NC strip, as Mean ± SD (n=3) and relative (in percentage) to the 500nM–T sample value. (·) denotes a significant difference when $p < 0.1$, compared to the negative control (NT); (*) when $p < 0.05$; (**) when $p < 0.01$; (***) when $p < 0.001$. Statistical analyses were performed by one-way ANOVA with Tukey's *post hoc* test.

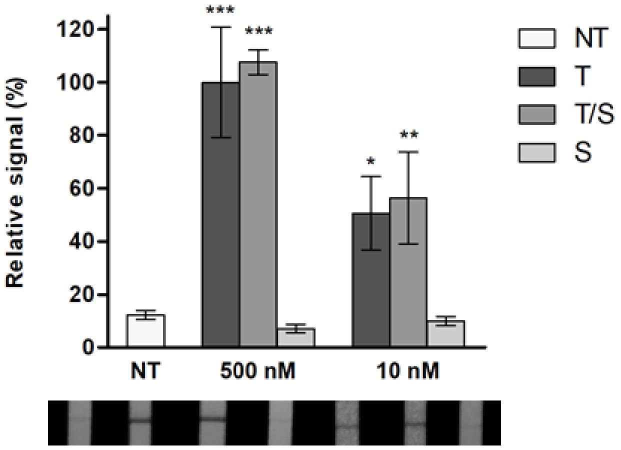

**Figure S9.** Multiplexed detection of the targets miR-146a, miR-155, and miR-191, in the presence of scramble sequences, using AuNP-Multi and streptavidin NC strips – photographs of the strips and signal intensity plots. From left to right were added: a negative control (NT – no target); a sample containing the three targets at the indicated total concentration (T); a mixture of the three targets and the six scramble sequences (S-1 to S-6), each one at the same concentration of each target (T/S); a sample containing only the six scramble sequences (S). The plots show the integrated signal for each NC strip, as Mean ± SD (n=3) and relative (in percentage) to the 500nM–T sample value. (*) denotes a significant difference when $p < 0.05$, compared to the negative control (NT); (**) when $p < 0.01$; (***) when $p < 0.001$. Statistical analyses were performed by one-way ANOVA with Tukey's *post hoc* test.

### 4. EXPAR amplification and detection of miRNAs

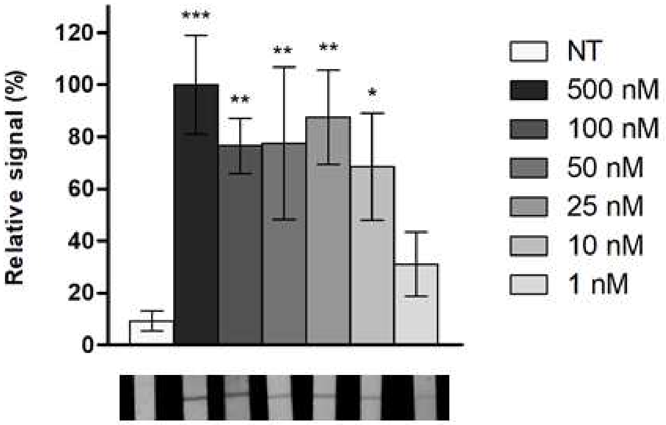

**Figure S10**. LoD determination of AuNP-EXP, using streptavidin NC strips – photographs of the strips and signal intensity plots. From left to right were added: a negative control (NT – no target); and the corresponding target sequence at different concentrations (from 500 nM to 1 nM – see legend). The plots show the integrated signal for each NC strip, as Mean ± SD (n=3) and relative (in percentage) to the highest intensity value (500 nM). (*) denotes a significant difference when $p < 0.05$, compared to the negative control (NT); (**) when $p < 0.01$; (***) when $p < 0.001$. Statistical analyses were performed by one-way ANOVA with Tukey's *post hoc* test.

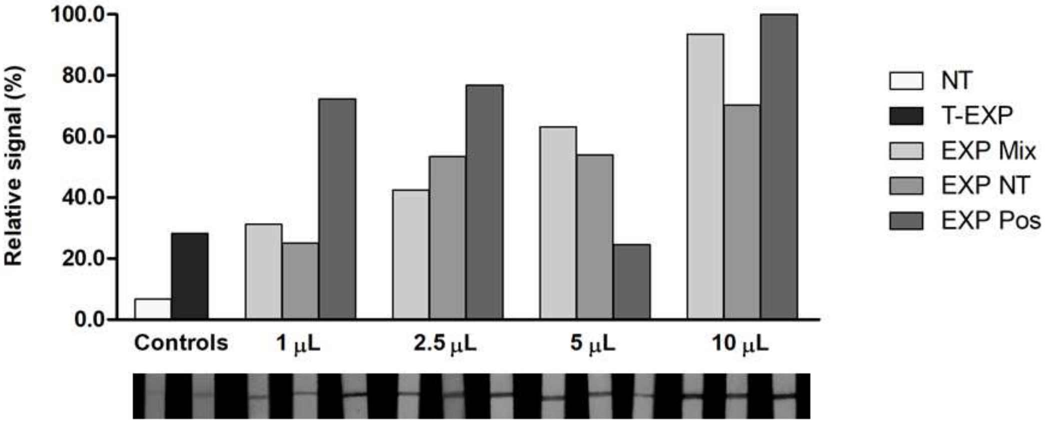

**Figure S11**. Detection of the T-EXP sequence in EXPAR reaction products, using AuNP-EXP and streptavidin NC strips (evaluation of the reaction volume to be incubated with the AuNPs) – photographs of the strips and signal intensity plots. From left to right were added: a negative control (NT – no target); the T-EXP synthetic sequence at 500 nM as a positive control (T- EXP); a no DNA EXPAR reaction mixture control (EXP-Mix); a negative EXPAR reaction (EXP-NT); and a mixture of positive EXPAR reactions (EXP-Pos). Four reaction volumes were tested: 1, 2.5, 5, and 10 µL. The plots show the integrated signal for each NC strip and relative (in percentage) to the highest intensity value (EXP-Pos, 10 µL).

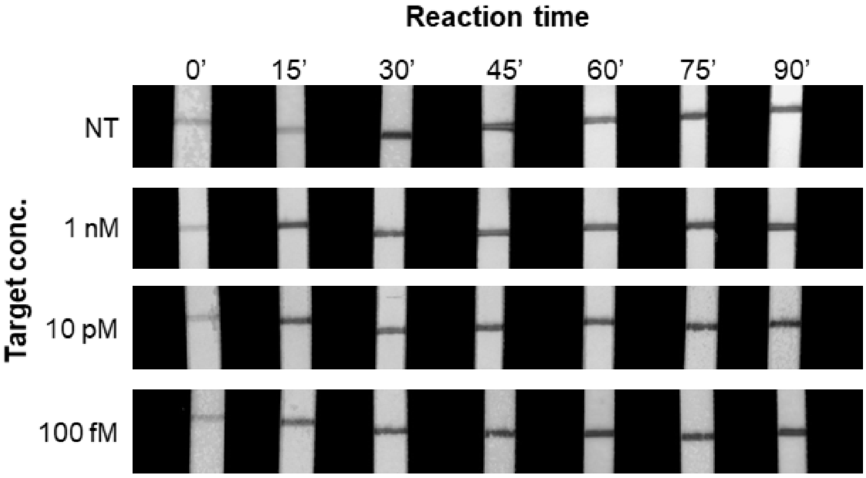

**Figure S12**. Detection of the T-EXP sequence in EXPAR reaction products, using AuNP-EXP and streptavidin NC strips (different reaction times) – photographs of the strips loaded with AuNP-EXP after incubation with the reaction products. Seven reaction times were evaluated: 0’, 15’, 30’, 45’, 60’, 75’, and 90’. The T-155 sequence was added at the concentrations of 1 nM, 10 pM, and 100 fM, compared to samples without target sequence (NT – no target).
